# An Innate Immune Receptor Toll-1 converts chronic light stress into glial-phagocytosis

**DOI:** 10.1101/2025.09.03.673940

**Authors:** Jiro Osaka, Toshiharu Ichinose, Mai Kanno, Satoko Hakeda-Suzuki, Takashi Suzuki, Atsushi Sugie

## Abstract

Chronic stress can cause progressive neuronal degeneration, yet the molecular mechanisms linking stress sensing to neuroimmune responses remain elusive. In this study, using a *Drosophila* model of chronic light-induced stress, we show that photoreceptor neurons accumulate reactive oxygen species (ROS) and exhibit Toll-1 activation which involves Spätzle ligands and receptor endocytosis. Toll-1 activation in neurons promotes axonal degeneration by inducing expression of the glial phagocytic receptor Draper (Drpr), leading to the engulfment of stressed axons. Genetic interaction analyses indicate that Toll-1 functions upstream of Drpr in a stress-responsive signaling cascade. Blocking either Toll-1 or Drpr attenuates axon loss under light stress, while Toll-1 overexpression exacerbates it. Toll-1 also plays a similar pro-degenerative role in an activity-dependent olfactory neuron degeneration paradigm, pointing to a broader role for this mechanism in neural degeneration. Together, these findings identify a neuron-glia signaling axis that converts sustained stress into structural degeneration.

## INTRODUCTION

Chronic stress has been increasingly recognized as a potent driver of progressive neuronal degeneration.^1,2^ Inflammatory responses often accompany this process within the nervous system, both in disease-associated and environmentally induced conditions,^3–6^ yet the underlying molecular mechanisms remain poorly understood. To address this, we previously established a chronic light exposure model in *Drosophila*, which recapitulates hallmarks of neurodegenerative conditions such as synapse loss and axon degeneration.^7^ Taking advantage of the fly’s stereotypic and spatially ordered visual system, we performed single-axon-resolution analyses using MeDUsA (<u>me</u>thod for the quantification of <u>d</u>egeneration <u>us</u>ing fly <u>a</u>xons), an automated tool we developed for quantifying photoreceptor axon loss in the optic lobe.^8^ Our model provides a valuable framework to investigate the molecular mechanisms by which chronic stress contributes to inflammatory neurodegeneration.

Toll-like receptors (TLRs) are conserved pattern recognition receptors that detect pathogen-associated molecular patterns (PAMPs) and initiate inflammatory responses.^9^ Prior to their discovery in mammals, the *Drosophila* Toll receptor was identified as essential for embryogenesis and later shown to mediate antifungal immunity, marking a breakthrough in innate immune research.^10^ Both *Drosophila* Tolls and mammalian TLRs share a similar structure—extracellular leucine-rich repeats (LRRs) and an intracellular TIR domain—suggesting a shared origin.^11^ Toll-1 is the primary mediator of immune defense in flies.^12^ The Toll-1 and immune deficiency (IMD) pathways activate NF-κB family factors (Dorsal and Relish) to induce antimicrobial peptide (AMP) genes, with Toll-1 mainly responding to Gram-positive bacteria and fungi, and IMD to Gram-negative bacteria.^13^ Other Toll family members, such as Toll-6 and Toll-7, are expressed in the nervous system and may function analogously to vertebrate neurotrophin receptors, regulating axon guidance and neuronal survival.^14,15^ In mammals, ten TLRs recognize various microbial components and trigger inflammation.^16^ Unlike mammals, Toll-1 is indirectly activated in flies: microbial recognition by PGRP-SA or GNBP1 leads to a serine protease cascade that activates the ligand Spätzle (Spz), which then binds Toll-1.^17^ Recent studies have shown that mammalian TLRs also detect endogenous danger signals (DAMPs), such as histones or protein aggregates released from damaged cells, inducing sterile inflammation.^18,19^ Similarly, non-infectious stress can activate immune pathways in *Drosophila*.^20,21^ For example, when apoptosis is blocked, intracellular contents can activate Toll-1 signaling via Persephone-mediated Spz cleavage.^22,23^ ROS produced by tissue damage also serve as triggers.^23–25^ Even minor mechanical injury can induce AMP expression through Toll-1 activation.^26^ Furthermore, stress signals from tumors can activate Toll-1 and IMD pathways, promoting immune responses that may suppress tumor growth.^27^ These findings suggest that the *Drosophila* Toll-1 pathway acts not only in pathogen defense but also as a sensor of internal stress, resembling mammalian sterile inflammation.

In addition to humoral immune responses such as antimicrobial peptide production, *Drosophila* also mounts a cellular immune response. In flies, glial cells and hemocytes clear apoptotic cells and foreign material, with the phagocytic receptor Draper (Drpr) playing a central role. Drpr, homologous to mammalian MEGF10 and MERTK, recognizes “eat-me” signals such as exposed phosphatidylserine and promotes debris clearance.^28,29^ Recent studies have shown that Toll-6 signaling enhances Drpr-mediated phagocytosis.^30^ In the larval CNS, apoptotic neurons release Spätzle 5 (Spz5), which activates Toll-6 on glial cells. This induces Drpr expression via the dSARM–FoxO pathway and enhances glial phagocytic capacity. Similar cooperation between Toll and Drpr has been observed in adult axonal injury models, suggesting their broader role in maintaining neural homeostasis.^31,32^ Loss of Drpr impairs clearance and leads to persistent inflammation and neurodegeneration.^33^ Conversely, excessive Drpr activity can also be harmful, leading to aberrant removal of healthy neurons and synapses.^33,34^ This imbalance forms a feedforward loop in which defective clearance promotes inflammation, further damages neurons.^35^ Under normal conditions, Drpr-mediated phagocytosis interrupts this loop and protects neural tissue. While Drpr maintains neural health by removing cell debris, excessive activity can damage the brain; for example, overexpression of *drpr* in glia leads to the removal of healthy neurons and synapses, resulting in neuronal loss, locomotor defects, and reduced longevity.^36,37^ These results indicate that Drpr activity must be tightly controlled. However, the mechanisms by which the nervous system senses and regulates glial phagocytic activity remain unclear.

In the present study, we utilized our established Drosophila model of chronic light stress-induced neurodegeneration to investigate how sustained stress promotes neuronal degeneration.^7,8^ We found that chronic light stress induces oxidative stress and mitochondrial dysfunction in photoreceptors. RNA-seq analysis revealed that light stress induces the expression of stress-responsive genes such as *X box binding protein-1* (*Xbp1*)*, Chaperonin containing TCP1 subunits* (*CCTs*) and *Heat-shock proteins* (*Hsps*), consistent with ROS accumulation. We further discovered that Toll-1 activation in neurons is a key driver of axon degeneration under chronic light stress. This degeneration is mediated by glial phagocytosis through the receptor Draper, revealing a previously unrecognized role for innate immune signaling in coordinating neuron–glia interactions during stress-induced neurodegeneration. These findings suggest that under chronic stress, neurons activate innate immune signaling to instruct glia to eliminate stressed axons, representing a unique mode of neuron-glia communication in sterile neurodegeneration.

## RESULTS

### Chronic Light Stress Induces ROS Accumulation and Alters Stress Response in the Photoreceptor

In our chronic light stress model, the photoreceptors undergo progressive neurodegeneration under continuous light (LL), but the underlying molecular mechanisms remain unknown (**Figure 1A–D**).^7^ Given previous studies that pigment cells protect photoreceptors by restricting excessive light exposure, we reduced pigmentation using the GMR-*white* RNAi transgene,^38,39^ which enabled us to recapitulate the slow-onset the axonal degeneration observed after 13 days of continuous light exposure, consistent with the observations in our earlier study (**Figure 1A–D**). Since previous studies have linked neuronal hyperactivity to ROS accumulation and neurodegeneration,^40–42^ we first tested whether ROS was elevated under LL. Using DCFH-DA, a non-fluorescent dye that is converted to the green-fluorescent compound DCF upon oxidation,^43^ we measured ROS in photoreceptors labeled with the red membrane marker mtdTomato.^44^ DCF signals were significantly higher under LL than under 12-hour light/dark cycle (LD) conditions, indicating light stress-induced ROS accumulation (**Figure 1E–G**). Furthermore, overexpression of ROS scavengers such as *sod1* and *cat* in the photoreceptor suppressed neurodegeneration, suggesting that ROS accumulation contributes directly to light-induced axonal loss (**Figure 1H**). We also observed mitochondrial fragmentation (**Figure S1A–D**), reduced autophagosomes (**Figure S1F–H**), and increased mitolysosomes upon chronic light exposure (**Figure S1J–L**). However, neither suppression of mitochondrial fragmentation via *drp1* RNAi (**Figure S1E**) nor genetic manipulation of autophagy- or mitophagy-related genes (**Figure S1I and S1M**) rescued neurodegeneration. These findings suggest that these changes are likely secondary consequences rather than direct drivers of degeneration.

**Figure 1.**
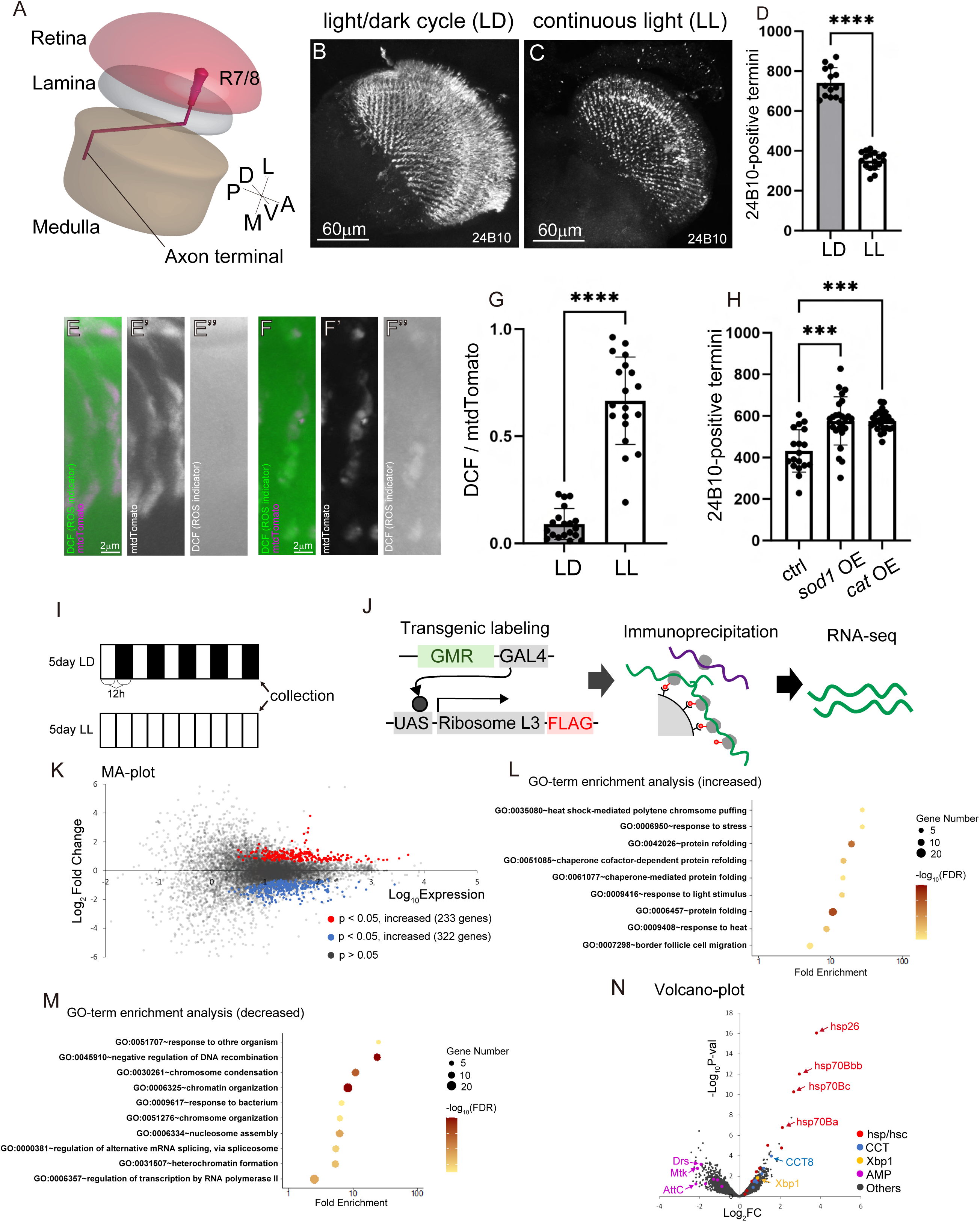
Molecular Changes in Photoreceptors under Light Stress. (A) A dorsal view schematic showing the organization of the *Drosophila* visual system, including the retina, lamina, and medulla. The axon terminals of photoreceptor neurons R7 and R8 are depicted projecting into their respective target layers within the medulla, exhibiting a stereotyped morphological pattern of axonal targeting. (B–D) Axons of R7 and R8 photoreceptors were visualized using mAb24B10 (white), a photoreceptor-specific antibody. 24B10-stained photoreceptor axons after 13 days of light/dark (LD) conditions and continuous light (LL) conditions. Scale bar, 60 μm. (D) Quantification of 24B10-positive photoreceptor axons under LD and LL conditions. Each point represents one brain; bars indicate mean ± SD. (E–F’’’) ROS signals detected by DCF in photoreceptors of flies maintained under LD or LL conditions for 5 days. Images show merged views (LD_Merge (E), LL_Merge (F)), photoreceptor axons (LD_Axons (E’), LL_Axons (F’)), and DCF signals (LD_DCF (E’’), LL_DCF (F’’)). Fly brains were dissected and immediately incubated with DCFH-DA solution, and DCF fluorescence was measured using confocal microscopy (see **Materials and Methods**). Scale bar, 2 μm. (G) Quantification of the volume ratio between DCF signals and photoreceptor axons (mtdTomato) of flies under 5 days of LD or LL. Each point represents one brain; bars indicate mean ± SD. (H) Quantification of 24B10-positive photoreceptor axons in control (40D-UAS), *sod1* overexpression (OE), and *cat* OE flies after 13 days of light stress. Each point represents one brain; bars indicate mean ± SD. (I) Light conditions used for TRAP-seq analysis. Photoreceptors from flies maintained under LD or LL for 5 days were collected. (J) Schematic diagram of the TRAP method. (K–N) Summary of TRAP-seq results and differentially expressed genes. MA plot (K), GO analysis of genes upregulated in LL (L), and GO analysis of genes downregulated in LL (M), Volcano plot (N). Statistical treatment of the number of 24B10-positive termini in (D) and volume ratio between DCF and mtdTomato in (G) were analyzed with the Mann-Whitney test. The number of 24B10-positive termini in (H) was statistically analyzed using the Kruskal–Wallis test with Dunn’s multiple comparisons test. The P-values in MA-plot (K) were calculated using DEseq2. In this and all following statistical significance. n.s. *P* > 0.05, * *P* ≤ 0.05, ** *P* ≤ 0.01, *** *P* ≤ 0.001, **** *P* ≤ 0.0001. See also **Figures S1** and **S2**.

To further investigate the molecular response to light stress, we performed Translating Ribosome Affinity Purification (TRAP)^45,46^ to isolate actively translated mRNAs from photoreceptors, followed by RNA-seq (**Figure 1I–N**). Flies expressing a FLAG-tagged ribosomal protein L3 (RpL3-FLAG) in the photoreceptor neurons were collected after 5 days of either LD or LL conditions (**Figure 1I and 1J**). mRNAs were immunoprecipitated, and successful enrichment of photoreceptor transcripts was confirmed by RT-PCR (**Figure S2**). RNA-seq was then performed (LD: n = 3; LL: n = 4), and differentially expressed genes (DEGs) were visualized using an MA-plot, identifying 232 upregulated and 322 downregulated genes under LL conditions (**Figure 1K**). Gene Ontology (GO) analysis showed enrichment of categories including heat shock response, protein folding, and response to heat/light stress among LL-upregulated genes (**Figure 1L**). In contrast, genes involved in immune responses, chromatin remodeling, and gene regulation were downregulated (**Figure 1M**). Volcano plot analysis revealed that stress-responsive genes such as *Xbp1*, *Hsp* family members, and *CCT* chaperonins were among the most significantly upregulated under LL conditions, whereas most antimicrobial peptide (AMP) genes showed downregulated expression (**Figure 1N**).^47–49^ Collectively, these data suggest that chronic light stress causes ROS accumulation in photoreceptor axons, leading to a compensatory upregulation of cellular stress response genes such as *Hsp* and *Xbp1*, while concurrently suppressing canonical inflammatory responses.

### Toll-1 is a Key Regulator of Photoreceptor Degeneration under Light Stress

RNA-seq analysis revealed that genes involved in stress responses, such as *hsp*, *xbp1*, and *cct*, were markedly upregulated under LL conditions (**Figure 1K–N**). In contrast, canonical inflammatory programs appeared to be downregulated, possibly to avoid additional toxicity under chronic stress. Previous studies have shown that such oxidative stress can activate the Toll-1 pathway.^23–25^ Therefore, we focused our subsequent analyses on this signaling axis. Consistent with a previous report,^50^ we confirmed the presence of Toll-1 in photoreceptor axons using a Toll-1::Venus knock-in line (**Figure 2A–A’’**).^51^ To test the functional relevance of Toll-1, we examined the function in our light-induced neurodegeneration model. In line with our earlier findings, pigment-deficient flies with a white mutant background without mini-white–containing transgenes exhibited progressive axonal degeneration within five days of continuous light exposure.^7^ Loss of *toll-1* or photoreceptor-specific knockdown exhibited significantly reduced axon degeneration, underscoring its essential and cell-autonomous role in the neurodegenerative process (**Figure 2B–E**). Conversely, photoreceptor-specific overexpression of *toll-1* accelerated axonal degeneration, even under early-stage light stress (**Figure 2F**). Together, these results demonstrate that Toll-1 is a critical mediator of light-induced axonal degeneration in the photoreceptor, functioning as a key link between environmental stress and structural decay.

**Figure. 2.**
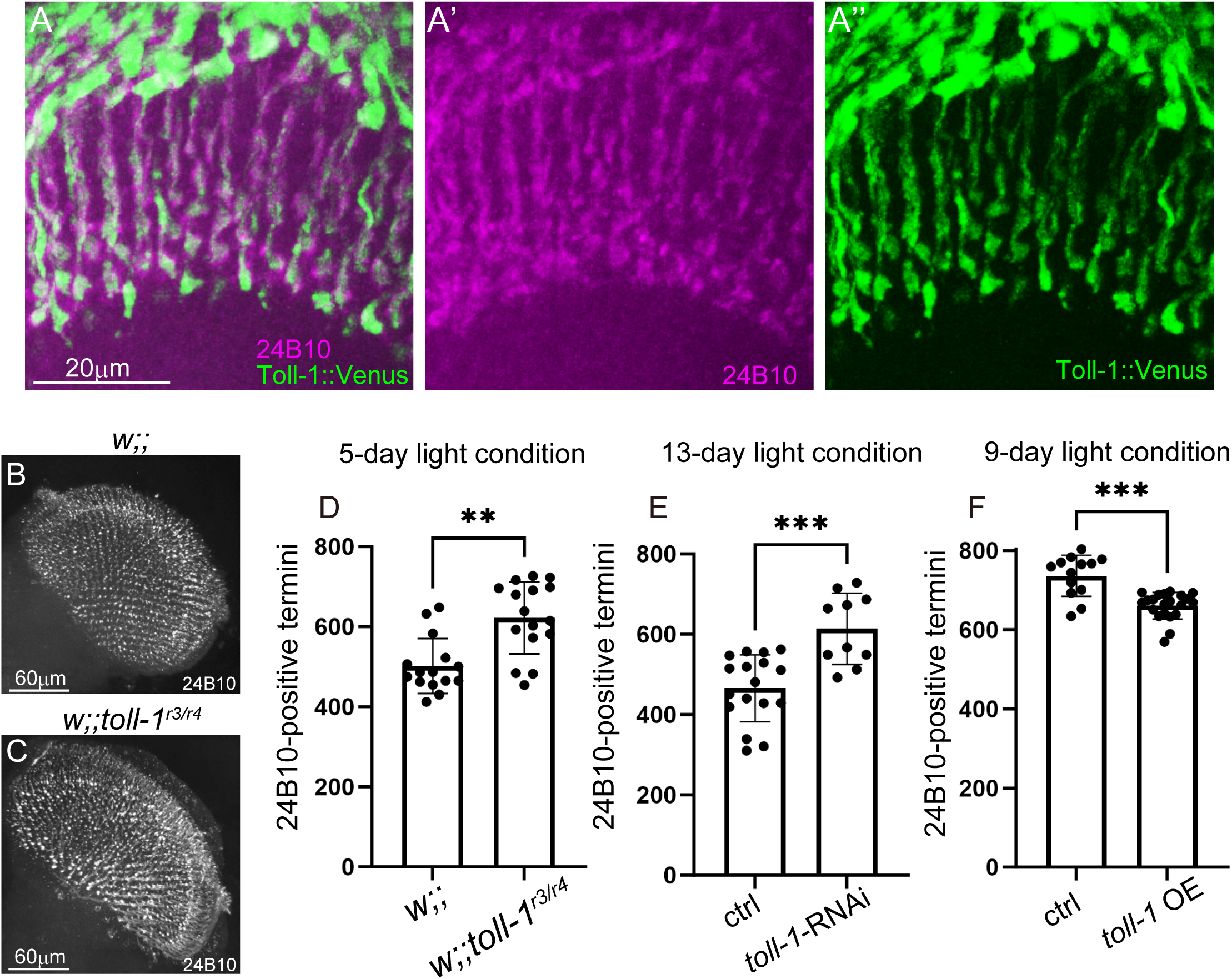
Toll-1 in Photoreceptors Mediates Light-induced Neurodegeneration. (A–A’’’) Endogenous localization of Toll-1. Shown are merged images (A), 24B10-stained photoreceptor axons (A’), and Venus-tagged Toll-1 knock-in signal (Toll-1::venus) (A’’). Scale bar, 20 μm. (B–D) Mutant analysis of *toll-1* for light-induced neurodegeneration. (B, C) 24B10-stained photoreceptor axons after 5 days of light stress in control flies (*w*;;) (B) and *toll-1* mutants (*w*;; *toll-1^r3/r4^*) (C). Scale bar, 60 μm. (D) Quantification of the 24B10-positive photoreceptor axons in control and *toll-1* mutant. Each point represents one brain; bars indicate mean ± SD. (E) Quantification of 24B10-positive photoreceptor axons in control (40D-UAS) and photoreceptor-specific *toll-1* RNAi flies after 13 days of light stress. Each point represents one brain; bars indicate mean ± SD. (F) Quantification of 24B10-positive photoreceptor axons in control (40D-UAS) and photoreceptor-specific *toll-1* overexpression (OE) flies after 9 days of light stress. *toll-1* OE induced premature axonal degeneration. Each point represents one brain; bars indicate mean ± SD. Statistical treatment of the number of 24B10-positive termini in (D–F) were analyzed using the Kruskal–Wallis test with Dunn’s multiple comparisons test.

### Endocytosed Toll-1 is Required for Light-induced Neurodegeneration

Previous studies have shown that Toll-1 endocytosis is important for its immune function in *Drosophila*.^52,53^ To investigate whether Toll-1 undergoes endocytosis in response to light stress, we examined the subcellular localization of Toll-1::Venus in photoreceptor axons. Under normal conditions, Toll-1::Venus was relatively uniformly distributed along the axons (**Figure 3A**). However, after exposure to continuous light, Toll-1::Venus formed punctate structures (**Figure 3B and 3C**). This formation of puncta was suppressed by blocking endocytosis using a temperature-sensitive dominant-negative form of dynamin, Shibire^ts1^ (**Figure 3D**).^54^ Moreover, Toll-1::Venus puncta colocalized with the early endosome marker Rab5 in photoreceptor axons (**Figure 3E–E’’**). Toll-1 also showed strong colocalization with endosomal markers such as Rab5 and Rab7 in *Drosophila* S2 cells (**Figure S3**), indicating that Toll-1 is capable of entering the endosomal pathway.

**Figure 3.**
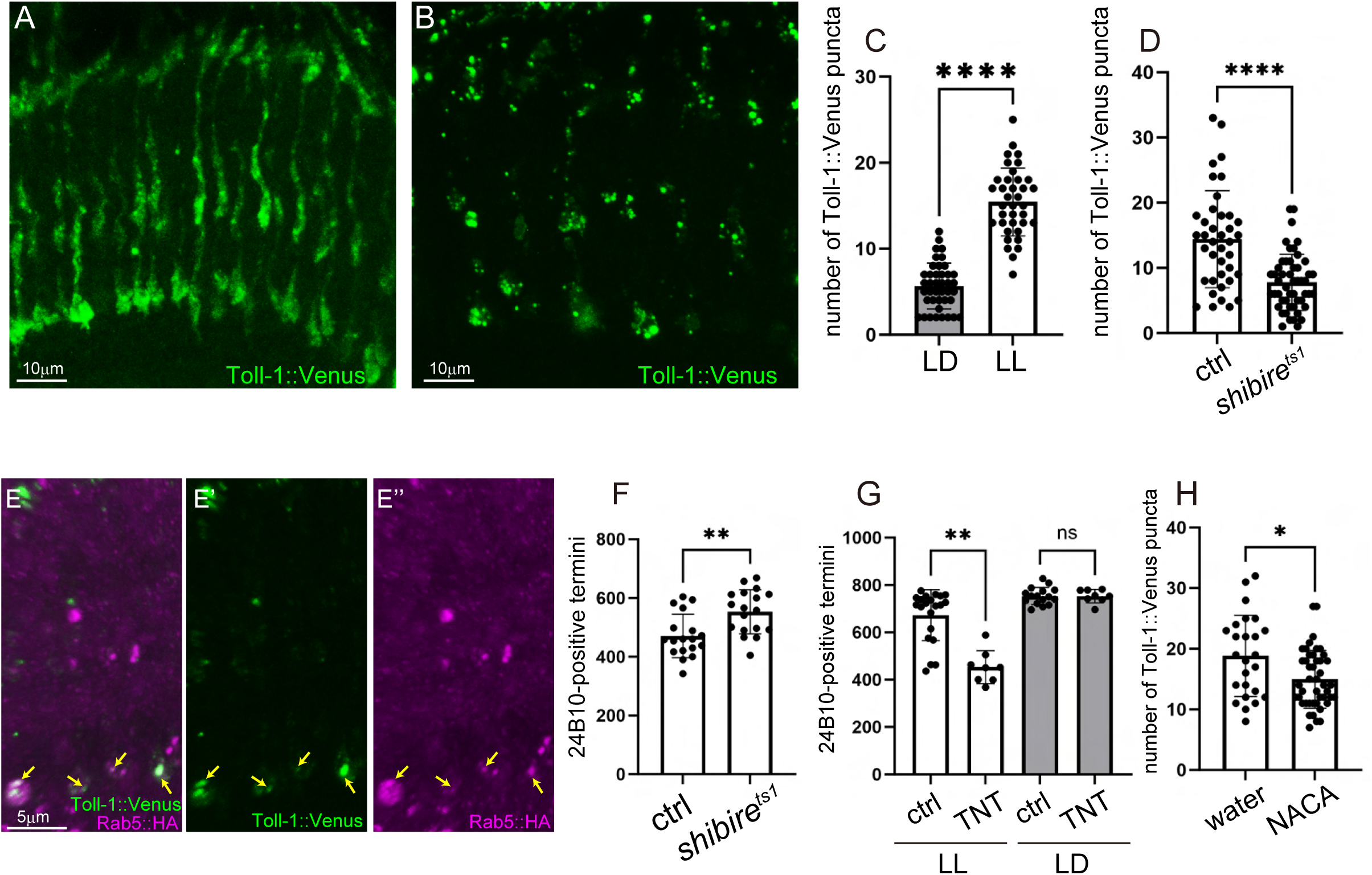
Toll-1 Internalized through Endocytosis under Light Stress is Involved in Neurodegeneration. (A–C) Localization of Toll-1::Venus in photoreceptors. *Toll-1::Venus* was expressed using the photoreceptor-specific driver GMR-GAL4. Images show flies kept under LD conditions for 5 days (A) and under LL for 5 days (B). Scale bar, 10 μm. (C) Quantification of Toll-1::Venus puncta per photoreceptor axon. Each point represents one axon; bars indicate mean ± SD. (D) Quantification of Toll-1::Venus puncta per photoreceptor axon with or without *shibire^ts1^* expressed by GMR-GAL4. Each point represents one axon; bars indicate mean ± SD. (E–E’’) Co-localization of Toll-1::Venus and the endosomal marker Rab5::HA in photoreceptors. Images show flies kept under LL conditions for 5 days. Merge (E), Toll::Venus (E’), and Rab5::HA (E’’). Arrowheads indicate puncta showing co-localization. Scale bar, 5 μm. (F and G) Analysis of the importance of endocytosis in light-induced neurodegeneration. Quantification of 24B10-positive photoreceptor axons in flies under LL for 13 days with endocytosis and synaptic transmission blocked by Shibire^ts1^ (F). Quantification in flies under LL or LD for 5 days with synaptic transmission blocked using 40D-UAS (control) and TNT, which specifically inhibits vesicle release. Each point represents one brain; bars indicate mean ± SD. (H) Quantification of Toll-1::Venus puncta per photoreceptor axon by feeding flies with water (control) or the antioxidant N-acetylcysteine amide (NACA). Each point represents one axon; bars indicate mean ± SD. Statistical treatment of the number of Toll-1::Venus puncta in (C, D, and H) and the number of 24B10-positive termini in (F) were analyzed with the Mann-Whitney test. The number of 24B10-positive termini in (G) was statistically analyzed using the Kruskal–Wallis test with Dunn’s multiple comparisons test. See also **Figures S3**.

Inhibition of endocytosis using Shibire^ts1^ significantly suppressed light-induced neurodegeneration (**Figure 3F**),^7^ indicating the importance of Toll-1 endocytosis. However, because Shibire^ts1^ also blocks synaptic vesicle recycling and neurotransmitter release,^54^ we tested whether neurotransmission itself was required for degeneration. Expression of the tetanus toxin light chain (TNT), which specifically inhibits vesicle release,^55^ failed to prevent axonal degeneration; instead, it markedly accelerated the process (**Figure 3G**). This finding underscores that the neuroprotective effect observed with Shi^ts1^ was due to the inhibition of endocytosis, rather than the cessation of synaptic vesicle recycling or neurotransmission itself. Furthermore, treatment with the antioxidant N-acetylcysteine amide (NACA)^56^ suppressed the light-induced formation of Toll-1 puncta in photoreceptor axons (**Figure 3H**). These findings suggest that light-induced oxidative stress promotes Toll-1 endocytosis in photoreceptors, which may play a role in initiating neurodegeneration.

### Spätzle Ligands Promote Toll-1–dependent Neurodegeneration

We next asked whether the light-induced neurodegeneration is mediated by known ligands of Toll-1, Spz and Spz5 (**Figure 4A**).^57,58^ To test this, we used a double mutant carrying null alleles of spz5 and spz (*spz5^Δ^*^44^*, spz^Δ8^*).^58^ In the pigment-deficient background used for this analysis, the double mutant showed significantly reduced axon degeneration following light stress, suggesting the role of Spz and/or Spz5 in this process (**Figure 4B-D**). We next examined whether overexpression of *spz* or *spz5* promotes light-induced neurodegeneration (**Figure 4E–I**). Under 9 days of light stress, which does not typically induce significant degeneration in control flies, *spz* overexpression showed a trend toward increased axon loss (p = 0.054), and *spz5* overexpression significantly enhanced degeneration (**Figure 4H**). Under more severe conditions (13-day light exposure), both *spz* and *spz5* overexpression robustly promoted axonal degeneration (**Figure 4E–G and I**).

**Figure. 4.**
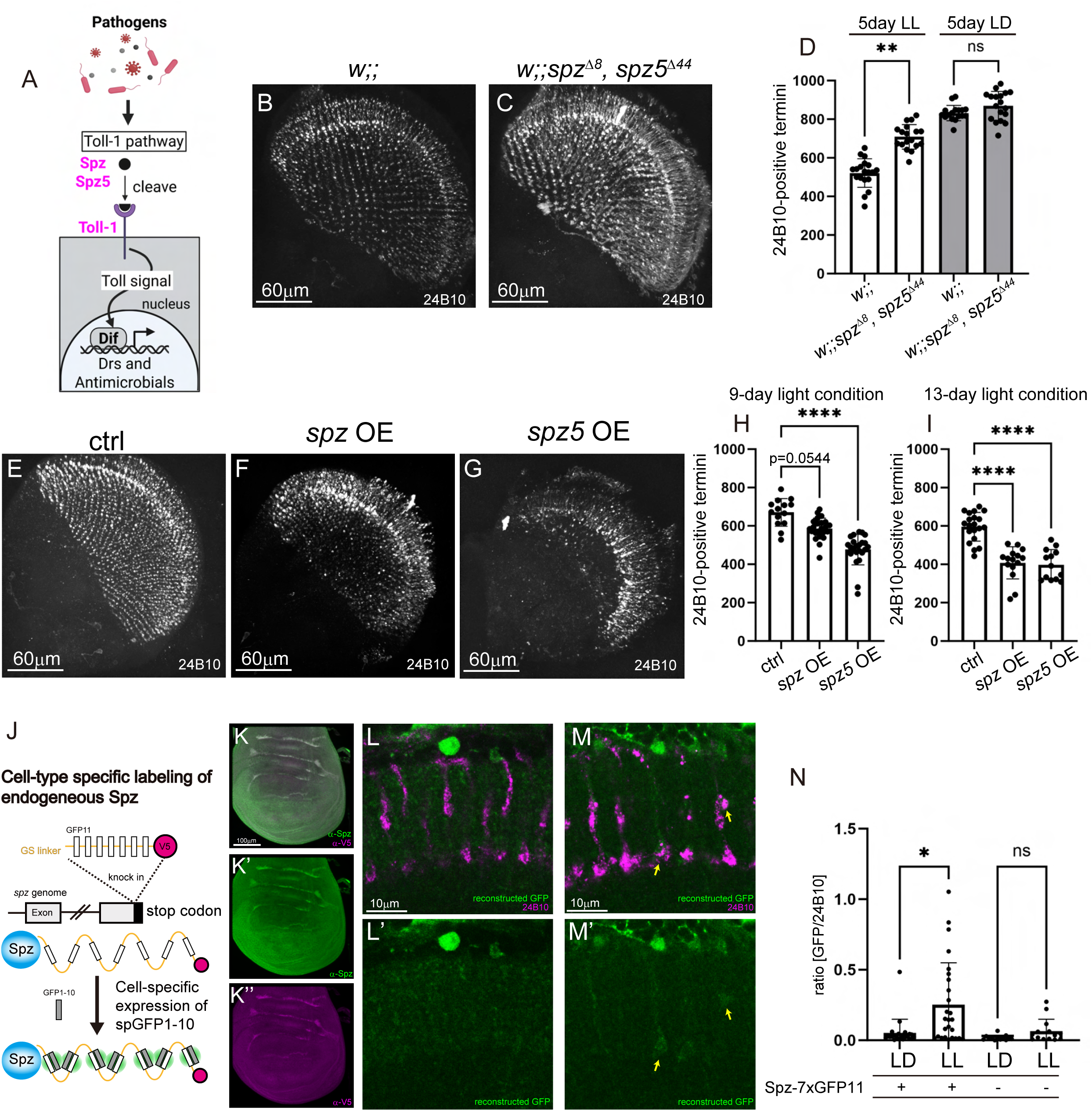
Spätzle Family Ligands Mediate Light Stress–Induced Neurodegeneration. (A) Schematic representation of the innate Toll-1 pathways in *Drosophila*. (B–D) Requirement of *spz* and *spz5* in light-induced neurodegeneration. (B, C) 24B10-stained photoreceptor axons after 5 days of LL. *w*;; (B), *w*;*;spz^Δ8^, spz5^Δ44^* (C). Scale bar, 60 μm. (D) Quantification of 24B10-positive photoreceptor axons in control (B) and the double mutant after 5 days of LL or LD (C). Each point represents one brain; bars indicate mean ± SD. (E-I) Overexpression of *spz* and *spz5* enhances light-induced neurodegeneration. 24B10-stained photoreceptor axons after 5 days of LL. ctrl (E), *spz* OE (F)*, spz5* (G). Scale bar, 60 μm. (H and I) Quantification of 24B10-positive photoreceptor axons in 40D-UAS (control) and photoreceptor-specific *spz* or *spz5* overexpression flies. Samples were exposed to light stress for 9 days (H) or 13 days (I). Each point represents one brain; bars indicate mean ± SD. (J) Schematic illustration of the strategy for cell-type-specific labeling of endogenous Spz. A tag containing seven tandem repeats of GFP11 linked by GS linkers together with a constitutive V5 tag was knocked into the endogenous *spz* locus. Cell-type-specific expression of spGFP1-10 enabled reconstitution of GFP fluorescence from endogenous Spz. (K–K’’) Validation of the endogenous Spz reporter in the wing disc. The constitutive V5 tag in the Spz knock-in reporter was detected by V5 immunostaining (K’) and showed strong colocalization with anti-Spz C106 antibody staining (K’’). Scale bar, 100 μm. (L–M’) Glial Spz around photoreceptor axons under LD (L and L’) and LL (M and M’). Endogenous Spz in glia was visualized by reconstituted GFP using repo-GAL4-driven spGFP1-10 expression, and photoreceptor axons were labeled with 24B10. Scale bar, 10 μm. (N) Quantification of the GFP/24B10 ratio under LD and LL conditions in flies with or without the Spz-7×spGFP11::V5 allele. Each point represents one brain; bars indicate mean ± SD. Statistical treatments of the number of 24B10-positive termini in (D, H, and I) and reconstructed GFP /24B10 (N) were statistically analyzed using the Kruskal–Wallis test with Dunn’s multiple comparisons test.

To visualize the localization and dynamics of endogenous Spz in specific cell types, we generated a split-GFP knock-in reporter in which Spz was tagged with a constitutive V5 tag and seven tandem GFP11 repeats, enabling Spz visualization in genetically-defined cells expressing the complementary GFP1-10 fragment (**Figure 4J**).^59^ We validated this reporter in the wing disc, where Spz localization has been well-characterized in previous studies.^60^ The V5 signal showed strong colocalization with anti-Spz antibody staining, indicating that the knock-in reporter faithfully recapitulates endogenous Spz localization (**Figure 4K–K’’**). Since previous RNA-seq data indicated the glial enrichment of *spz* expression in the adult brain,^61^ we analysed glial Spz using the pan-glial driver repo-GAL4. We found that glial Spz level surrounding photoreceptor axons were increased under LL conditions compared with LD conditions (**Figure 4L–N**). Together, these findings suggest that Spz, upregulated in glial cells under chronic light stress, promote light-induced neurodegeneration, likely through Toll-1 signaling.

### Glial Draper-mediated Phagocytosis is Essential for Light-induced Neurodegeneration

Since our analysis revealed that Toll-1 and its ligands promote axonal degeneration, we next investigated the potential involvement of glial phagocytosis, a process previously shown to be regulated by Toll-6 signaling during development.^30^ We tested whether a similar mechanism operates under chronic light stress. In *Drosophila*, the key glial phagocytic receptor is Drpr.^62^ Under LD conditions, GFP-knock-in Drpr signals near the photoreceptor were faint (**Figure 5A–A’’**). In contrast, LL conditions led to a marked increase in Drpr signal around photoreceptor axons, with approximately 80% of photoreceptor axons wrapped by the Drpr signal (**Figure 5B–C**). In addition, downstream effectors of Drpr signaling—Stat92E and AP-1—were both upregulated in response to light stress (**Figure S4**), supporting activation of glial phagocytic pathways. To directly test whether glial phagocytosis is required for neurodegeneration, we next examined a *drpr* mutant in a pigment-deficient background. Axon degeneration was strongly suppressed in these mutants under LL, with almost no axon loss observed (**Figure 5D–F**). This striking rescue, combined with the morphological analyses (**Figure 5A-C)**, indicates that glial phagocytosis is essential for light-induced degeneration. Consistent with this, pan-neuronal knockdown of *drpr* did not suppress neurodegeneration, whereas glial-specific knockdown of *drpr* did (**Figure 5G–I**), confirming that Drpr function in glia is required for axon loss. To explore whether axons externalize phosphatidylserine (PS), an “eat-me” signal that acts as a ligand for Drpr,^63,64^ we employed the PS reporter GFP-LactC1C2^65^ secreted from nearby cells (Dm8, L2, and glia). Using this reporter, we found little to no externalize PS signal on axons under LD conditions, but strong labeling under LL (**Figure 5J–P**). Together, these findings suggest that Toll-1–induced, Drpr-mediated phagocytosis by glial cells, likely facilitated by the recognition of exposed PS, drives axonal degeneration under chronic light stress.

**Figure. 5.**
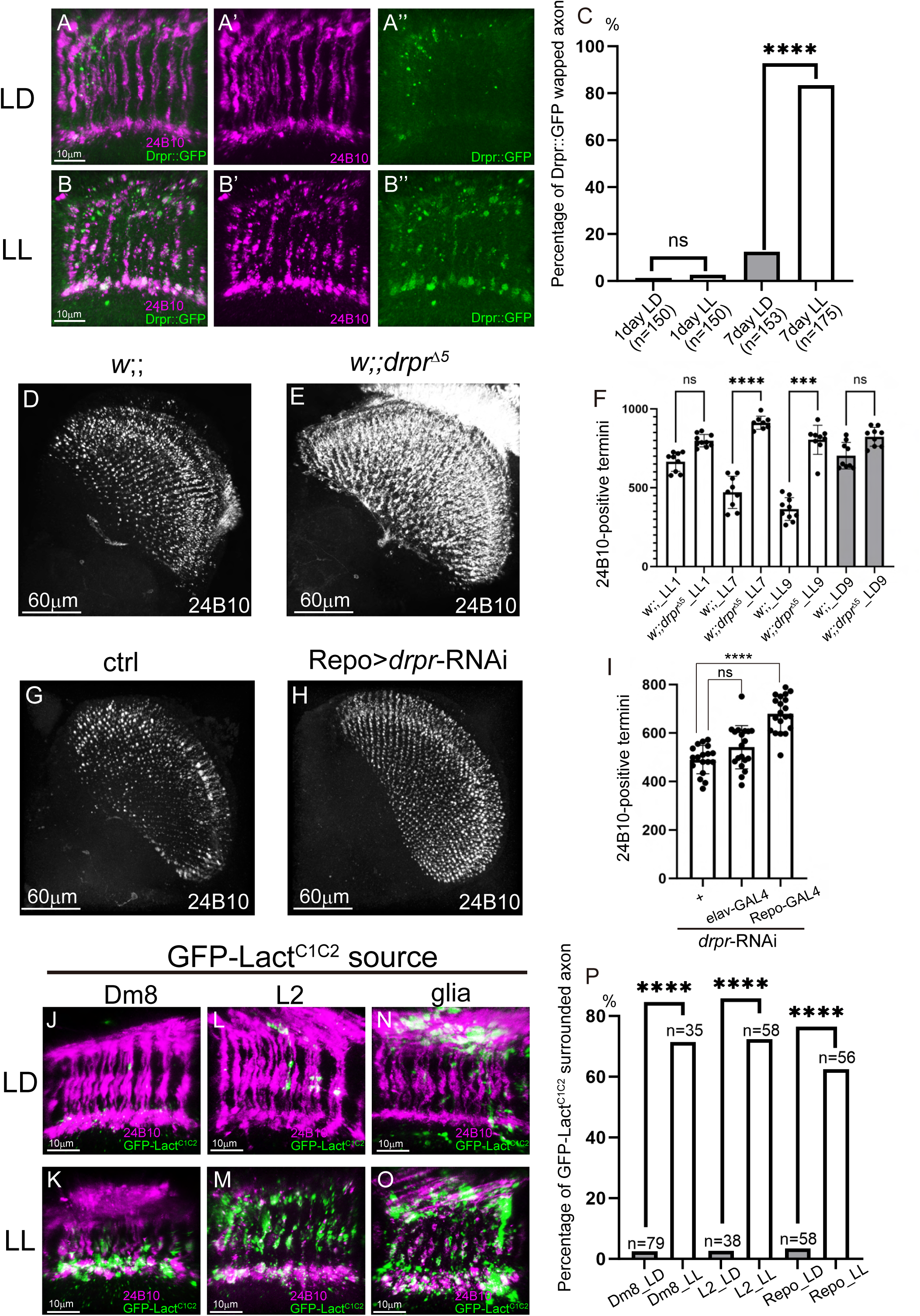
Excessive Neuronal Clearance by Drpr Underlies Light-Induced Photoreceptor Axon Degeneration. (A–B’’) Endogenous localization of Drpr with Drpr::MiMIC-GFP transgenic flies under different light conditions. Photoreceptors from flies exposed to 7 days of LD (A–A’’) and 7 days of LL (B–B’’). Scale bar, 10 μm. (C) Quantification of the proportion of photoreceptor axons sFurrounded by Drpr signal under LD and LL conditions at day 1 and day 7. (D–F) Requirement of *drpr* for light-induced neurodegeneration. Mutant analysis of *drpr* for light-induced neurodegeneration. (D and E) 24B10-stained photoreceptor axons after 9 days of LL in control flies (*w*;;) (D) and *drpr* mutants (*w*;; *drpr^Δ5^*) (E). Scale bar, 60 μm. (F) Quantification of 24B10-positive axons under several light conditions. Each point represents one brain; bars indicate mean ± SD. (G–I) Requirement of *drpr* for light-induced neurodegeneration in glial cell. 24B10-stained photoreceptor axons after 13 days of LL in control flies (G) and glial *drpr* RNAi flies (H). Scale bar, 60 μm. (I) Quantification of 24B10-positive axons in control and neural or glial *drpr* RNAi under 13 days of LL. Each point represents one brain; bars indicate mean ± SD. (J–O) Detection of phosphatidylserine exposure using the secreted GFP-Lact^C1C2^ marker expressed from surrounding cells under 7 days of LD or LL. (J and K) GFP-Lact^C1C2^ expressed from Dm8 neurons under LD (J) and LL (K) conditions. (L and M) GFP-Lact^C1C2^ expressed from L2 neurons under LD (L) and LL (M) conditions. (N and O) GFP-Lact^C1C2^ expressed from glial cells under LD (N) and LL (O) conditions. Scale bar, 10 μm. (P) Quantification of the proportion of photoreceptor axons surrounded by GFP-Lact^C1C2^ signal under 7 days of LD or LL. Statistical treatment of the percentage the Drpr::GFP wrapped in (C) or GFP-Lact^C1C2^ surrounded axon in (P) were analyzed with Fisher’s Exact test. Ther number of 24B10-positive termini in (F) and (I) were statistically analyzed using the Kruskal–Wallis test with Dunn’s multiple comparisons test. See also **Figure S4**.

### Drpr-mediated Phagocytosis is Promoted Downstream of Toll-1

To clarify how Toll-1 contributes to axonal degeneration under chronic light stress, we examined its relationship with the glial phagocytic receptor Drpr. We first asked whether Toll-1 activity affects the induction of Drpr expression after light stress. In wild-type flies, Drpr signals were observed in approximately 80% of photoreceptor axons following LL exposure. In contrast, this proportion dropped to nearly half in *toll-1* heterozygotes (**Figure 6A–C**), suggesting that Toll-1 is required for full activation of Drpr-mediated phagocytic responses. To further test whether Toll-1 and Drpr act in the same genetic pathway, we performed a double heterozygote analysis (**Figure 6D–H**). Neither heterozygous mutation of *toll-1* nor *drpr* alone significantly suppressed neurodegeneration (**Figure 6D–F**), but the combination of both heterozygous mutations led to a strong reduction in axon loss (**Figure 6G and H**). These findings indicate a genetic interaction between Toll-1 and Drpr and suggest that Toll-1 promotes glial engulfment of stressed axons by acting upstream of Drpr.

**Figure 6.**
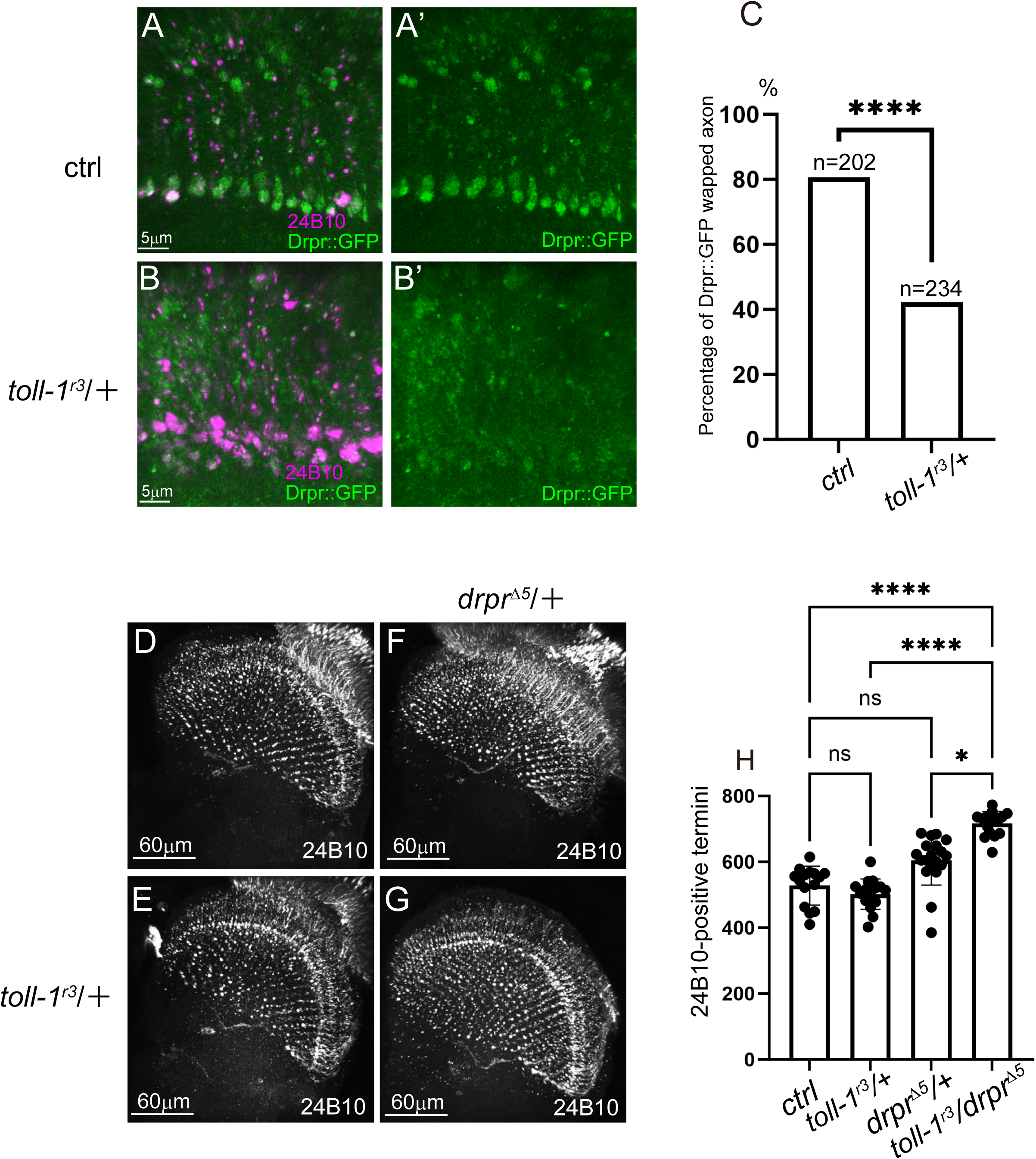
Toll-1 Functions upstream of Drpr to Promote Glial Phagocytosis under Chronic Light Stress. (A–B’) Endogenous Drpr signals in photoreceptor axons after 7 days of LL. Control (A and A’) and *toll-1* heterozygous mutant (B and B’). Scale bar, 5 μm. (C) Quantification of the proportion of photoreceptor axons wrapped by Drpr signal after 7 days of LL. *toll-1* heterozygous mutants exhibited a significant reduction in Drpr-positive axons compared to controls. (D–H) Genetic interaction analysis between *toll-1* and *drpr*. (D–G) 24B10-stained photoreceptor axons after 8 days of light stress in control (D), *toll-1* heterozygous mutant (E), *drpr* heterozygous mutant (F), and *toll-1* and *drpr* heterozygous mutant (G). Scale bar, 60 μm. (H) Quantification of 24B10-positive axons in 8 day light condition. All flies used in this assay carried GMR-*white*-RNAi and GMR-*dicer2* transgenes on the first chromosome, which drastically reduce eye pigmentation. As a result, even control flies exhibited photoreceptor degeneration after 8 days of light stress. Each point represents one brain; bars indicate mean ± SD. Statistical treatment of the percentage the Drpr::GFP wrapped axon in (C) was analyzed with the Chi-square test. Ther number of 24B10-positive termini in (H) was statistically analyzed using the Kruskal–Wallis test with Dunn’s multiple comparisons test.

### Toll Receptors Regulate Activity-Dependent Neurodegeneration in Olfactory Neurons

To determine whether the Toll-driven neuroinflammation–phagocytosis mechanism observed in the visual system generalizes to other sensory modalities, we employed an olfactory degeneration paradigm in *Drosophila* (**Figure 7A**). In this model, chronic exposure to ethyl butyrate (EB) induces the degeneration of Or42a olfactory receptor neurons (ORNs), a process previously reported to involve glial-mediated phagocytosis.^66–70^

**Figure 7.**
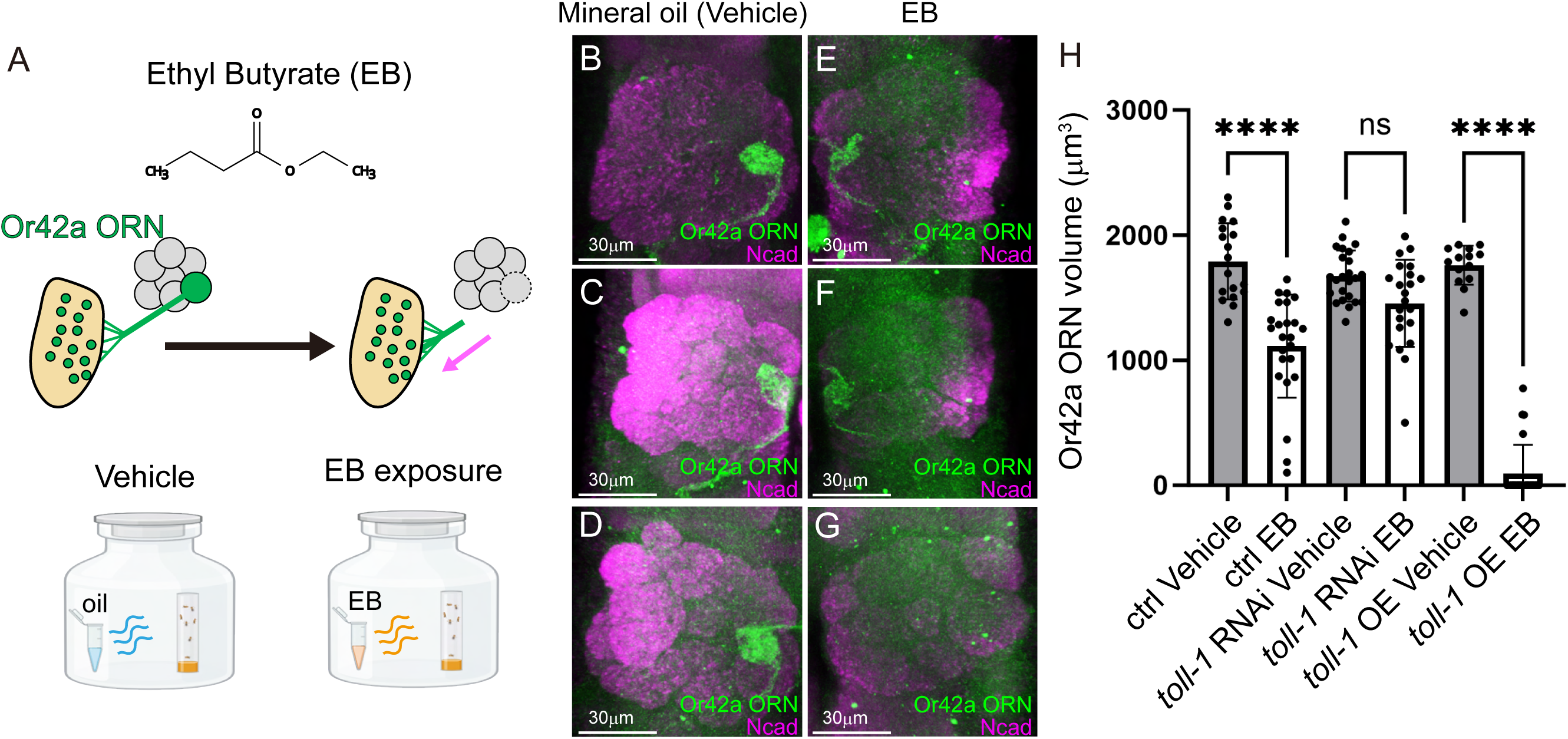
A Conserved Pro-degenerative Role of Toll-1 in the Or42a Olfactory Receptor Neuron Model. (A) Schematic representation of the experimental design. Top: Anatomy of the adult *Drosophila* maxillary palp, showing projections of Or42a ORNs (green) to the corresponding glomerulus (green circle). Activity-dependent degeneration induced by EB, an odorant that strongly activates Or42a ORNs, induces neuronal degeneration. Bottom: For odor stimulation, flies were placed in a glass chamber containing either vehicle (mineral oil) or 15% EB, and were analyzed after 2 days of exposure. (B–G) Axon of Or42a ORN under Mineral oil (vehicle) condition (B–D) or 2 days of exposure to 15% EB (E–G). Or42a ORNs were visualized by Or42a-GAL4-driven UAS-mCD8::GFP (green), and neuropil was labeled with N-cadherin (Ncad) (magenta). ctrl (40D-UAS) (B and E), *toll-1* RNAi (C and F), and *toll-1* overexpression (OE) (D and G). Scale bar, 30 μm. (H) Quantification of Or42a ORN axon volume per glomeruli under vehicle conditions and 2day 15% EB condition. Each point represents one brain; bars indicate mean ± SD. Statistical treatment the volume of Or42a ORN axons was statistically analyzed using the Kruskal–Wallis test with Dunn’s multiple comparisons test.

We first examined the morphology of Or42a ORNs under vehicle (mineral oil) conditions. In the absence of EB, the glomerular structures remained intact across all genotypes, including the control (**Figure 7B**), *toll RNAi* (**Figure 7C**), and *toll OE* (**Figure 7D**) flies. Upon EB exposure, control animals exhibited significant regression of Or42a ORN projections (**Figure 7E**). We found that knockdown of *toll-1* markedly suppressed this EB-induced degeneration (**Figure 7F**). Conversely, overexpressing *toll-1* dramatically exacerbated the phenotype, leading to a near-complete loss of glomerular innervation following EB exposure (**Figure 7G**). Quantitative analysis of Or42a ORN volume confirmed these observations (**Figure 7H**). Collectively, these data demonstrate that the Toll–Drpr axis is a conserved molecular link between neuronal activity and glia-mediated neurodegeneration, functioning in both the visual and olfactory systems.

## DISCUSSION

### Toll Receptor Endocytosis and Spz-Dependent Signaling Mediate Light-Induced Neurodegeneration

In this study, we show that photoreceptor axon-loss, mediated by Toll-1 activation, requires endocytosis. Upon light-induced stress, Toll-1 is internalized and localizes to endosomes (**Figure 3A–D**). Blocking endocytosis with Shibire^ts1^ mutant suppressed both Toll-1 endocytosis and light-induced neurodegeneration, suggesting that endocytosis is a key step in Toll-1 signaling leading to neurodegeneration (**Figure 3F**). Previous studies have also suggested that endocytosis plays an important role in Toll-1 activation,^52,53^ and our findings support this idea. These findings raise the possibility that internalized Toll-1 contributes to PS externalization, potentially facilitating the Draper-mediated glial phagocytosis observed in this model. Moreover, the constitutively active Toll-1^10b^ allele has been reported to increase endocytosis, implying that Toll-1 activation may depend on Spz binding and subsequent internalization.^53^ This raises the possibility that, following ligand binding at the cell surface, Toll-1 may be internalized together with Spz and continue signaling from endosomal compartments. We found that Spz and Spz5 function as critical ligands that activate Toll-1 and promote neurodegeneration. Flies carrying null alleles both *spz* and *spz5* showed a marked reduction in degeneration (**Figure 4B–D**). In contrast, when either *spz* or *spz5* was overexpressed, axon loss was accelerated (**Figure 4E–I**). These findings indicate that Spz/Spz5-dependent activation of Toll-1 contributes to neurodegeneration in the adult CNS. A similar mechanism has been described during development, where apoptotic neurons release Spz5, which activates Toll-6 in glia to induce Draper expression and promote phagocytosis.^30^ Our findings suggest that the adult Toll-1–Spz pathway may represent a parallel mechanism for regulating glial phagocytosis. Importantly, while Toll-1 functions in photoreceptors, other Toll receptors have been reported to be expressed in different parts of the nervous system and play roles in distinct neural contexts.^14,15,50,61,71,72^ This implies that multiple Toll receptors could regulate glial clearance in the adult brain, depending on the cellular context. Together, our results reveal a previously unrecognized pathway in which endocytosis and Spz ligands activate Toll-1 in photoreceptors to trigger neurodegeneration through glial phagocytosis. This mechanism may represent an adult-specific variant of a developmental program, with distinct receptors and modes of activation.

### Light-Induced ROS Accumulation as a Candidate Driver of Sterile Toll-1 Pathway Activation

Because this model lacks microbial infection, Toll-1 pathway activation may be mediated by cleavage of Spz ligands in response to cellular stress. Light stress leads to increased levels of ROS (**Figure 1E–G**), and the neurodegenerative phenotype can be rescued by ROS scavengers (**Figure 1H**), suggesting that ROS acts upstream of Toll-1 signaling. Structural abnormalities in mitochondria (**Figure S1A–D**), a major source of ROS, were observed under light stress. Consistently, mito-QC analysis indicated impaired mitochondrial quality (**Figure S1J–L**). Furthermore, feeding flies with the antioxidant NACA suppressed light-induced Toll-1 activation **(Figure 3H**). These findings suggest that mitochondrial dysfunction under light stress leads to increased ROS production, which may act as a trigger for Spz processing and subsequent Toll-1 activation, ultimately promoting neurodegeneration. Previous studies have shown that ROS activates serine protease cascades, which cleave Spz precursors to initiate Toll-1 signaling.^23^ In a sterile injury model in adult *Drosophila*, hydrogen peroxide (H₂O₂) produced by the enzyme Duox triggers activation of the protease Persephone (Psh), which in turn cleaves Spz, which is required to activate the Toll-1 pathway.^23^ These findings suggest that chronic light exposure leads to Toll-1 activation through ROS accumulation and Spz ligands cleavage even in the absence of infection or tissue injury. Therefore, ROS may function as an endogenous DAMP, activating Toll-1 signaling through Spz processing, although direct evidence remains to be established. Finally, how secreted Spz and Spz5 precursors are locally cleaved under light stress remains unresolved. Identifying the specific serine proteases responsible for this process is an important direction for future research.

### Glial Phagocytosis via the Toll–Drpr Axis Drives Neurodegeneration

This study demonstrates that activation of the Toll promotes the expression and function of the glial phagocytic receptor Drpr, leading to neurodegeneration. Downregulation of *toll-1* abolished light stress–induced upregulation of Drpr (**Figure 6A–C**), and *drpr* mutants failed to exhibit neurodegeneration under light stress conditions (**Figure 5D–F**). These findings suggest that the Toll–Drpr axis contributes to photoreceptor loss by promoting glial phagocytosis in a non–cell-autonomous manner. In this context, photoreceptors are not eliminated solely through intrinsic cell death pathways triggered by stress; rather, they are actively removed by surrounding glial cells through Toll-dependent phagocytic activity. This mechanism is consistent with the concept of “phagoptosis,” in which otherwise viable neurons are selectively targeted and cleared by glial phagocytes. It also parallels a previously reported developmental pathway, in which neuronal Spz5 activates glial Toll-6 to induce Drpr expression and promote selective neuronal clearance,^30^ suggesting that a similar mechanism operates in the adult nervous system. Consistent with this idea, Toll-1 also promoted degeneration in an EB-induced Or42a ORN degeneration paradigm (**Figure 7**), a distinct glial phagocytosis-dependent model, suggesting that this role may be conserved across neural contexts. Importantly, Drpr-mediated phagocytosis is not always pathological. In flies lacking Drpr, undegraded apoptotic cells accumulate, leading to age-related brain degeneration and reduced lifespan.^33,34^ This highlights that glial phagocytosis is essential for tissue homeostasis and that both insufficient and excessive activity can be harmful. Other studies also support non–cell-autonomous mechanisms of degeneration under light stress. For instance, in *norpA* mutants, sustained light exposure causes Rh1 accumulation, which activates the immune Imd–Relish pathway and induces cell death.^73^ Another model suggests that excess neuronal activity increases ROS levels, leading to lipid droplet accumulation in glia, which contributes to neurodegeneration.^74^ These examples align with our findings, emphasizing the role of glial responses in stress-induced neuronal damage. Moreover, in mammalian neurodegenerative diseases, glial phagocytosis and inflammation have dual roles. In early Alzheimer’s disease and glaucoma, overactive microglia aberrantly remove healthy synapses and neurons,^75^ mirroring the Toll–Drpr–dependent clearance of live neurons described here. Conversely, moderate activation via TREM2 has been shown to support Aβ clearance and promote tissue repair,^76^ indicating that the quality and timing of glial responses critically influence disease outcomes.

In conclusion, this study identifies Toll-mediated glial phagocytosis as a non–cell-autonomous mechanism of neurodegeneration. Disruption of this regulatory axis may initiate pathological neurodegeneration. Future studies should investigate whether a similar cascade—stress-induced release of endogenous ligands, activation of TLR or IL-1 pathways, microglial activation and phagocytosis, and neuronal loss—also operates in mammalian systems. Our findings suggest that the Toll–Drpr axis may serve as a critical regulator in the balance between protective and pathological glial responses and offer a potential target for therapeutic intervention.

### Potential Involvement of a Novel Toll-1 Pathway in Neurodegeneration

Our RNA-seq analysis revealed that while stress-response genes encoding heat shock proteins were strongly induced under chronic light stress, expression of AMP genes like *drs* decreased. Gene Ontology categories such as “response to bacterium” and “response to other organism” were also significantly enriched in the downregulated genes (**Figure 1M**). These findings suggest that canonical Toll-1 immune responses such as AMP production are selectively suppressed under our sterile environmental stress conditions. This may reflect the absence of pathogenic infection in our model and implies context-dependent modulation of the Toll-1 pathway. Consistently, inhibiting canonical downstream components of the Toll-1 pathway—including the Myd88-dependent pathway,^77,78^ the TRIF/TRAM pathway (Myd88-independent),^79,80^ and the Wek/Sarm/JNK pathway^81,82^—did not produce similar protective effects in our model (**Figure S5A–D**). Moreover, transcript levels of these downstream factors were not elevated in our TRAP-seq data (**Figure S5E**). This pattern may reflect immune tolerance or negative feedback mechanisms that suppress immune gene expression during prolonged stimulation or chronic stress.^83^ Alternatively, the observed upregulation of Hsp-related genes (**Figure 1N**) suggests a shift in cellular resources from AMP production toward other stress-response pathways such as Hsp synthesis.^84,85^

Taken together, our results indicate that canonical Toll-1 downstream pathways are unlikely to mediate neurodegeneration in this model. Instead, Toll-1 may engage a previously unidentified signaling branch under chronic light stress. This study highlights the possibility that Toll-1 functions independently of classical AMP induction and may act via a novel, stress-responsive neurodegenerative pathway. Identifying this pathway will be crucial for understanding how chronic stress contributes to neurodegeneration.

### Translational Relevance and Species-Specific Considerations

This study is based on a *Drosophila* model, and care should be taken when applying these findings to mammals. Toll-like receptors vary across species: in flies, Toll-1 is primarily activated by the ligand Spz, whereas in mammals, TLRs directly recognize pathogen- or damage-associated molecules. While the upstream activation mechanisms differ, downstream components such as NF-κB signaling and phagocytic pathways are highly conserved. Additionally, the mammalian nervous system is far more complex, involving diverse glial populations and multiple overlapping immune and inflammatory pathways. As a result, the effects of manipulating a single pathway may be less pronounced or more context-dependent compared to the fly model. Nonetheless, the core mechanism proposed in this study—stress-induced release of endogenous ligands driving glial phagocytosis and subsequent neurodegeneration—offers a valuable framework for exploring similar processes in mammals.

## Supporting information

Key Resource Table

## ACKNOWLEDGMENTS

We gratefully acknowledge Daiki Umetsu at Osaka University, Takayuki Kuraishi at Kanazawa university, Chun Han at Cornell University, Marc Freeman at Vollum Institute, and Masayuki Miura at National Institute for Basic Biology for providing the transgenic fly reagents. We thank Yuichi Shichino (RIKEN) for technical inputs regarding RNA-seq analysis. We thank Miki Yamamoto-Hino and Satoshi Goto at Rikkyo University for providing the anti-Spz C106 polyclonal antibody. We thank the BDSC, VDRC, and KYOTO Drosophila Stock Center for providing fly stocks. We thank BioRender for making Graphical abstract. This work was supported by JSPS KAKENHI (# 23K19651, 24KJ0089, and 24K19784 to J. O.; 25K22463 to T.I.; 24K09467 to S.H.S; #21H05682, #21H02483, #23H04220, 25K02281, and 25K22466 to T.S; 24K02349 and 24K22104 to A.S;), a Japan Agency for Medical Research and Development (#24ek0109760s401 (A.), a Takeda Science Foundation (T.S.; A. S.). Lotte Foundation (T.I.). Computations on the RNA-seq data were performed on the supercomputer at ROIS National Institute of Genetics.

## AUTHOR CONTRIBUTIONS

J.O., A.S., and T. I. designed experiments. J.O., A.S., T. I. and M.K. performed experiments and analysed data. J.O. and A.S. generated transgenic reagents. S.H.S. and T. S. provided experimental reagents and support. J.O., A.S. and T. I. wrote the manuscript with review from all co-authors.

## DECLARATION OF GENERATIVE AI AND AI-ASSISTED TECHNOLOGIES IN THE WRITING PROCESS

The authors used ChatGPT to improve the manuscript. After using ChatGPT, the authors reviewed and edited the content as needed and take full responsibility for the content of the publication.

## LEGENDS TO SUPPLEMENTAL FIGURES

**Figure S1.**
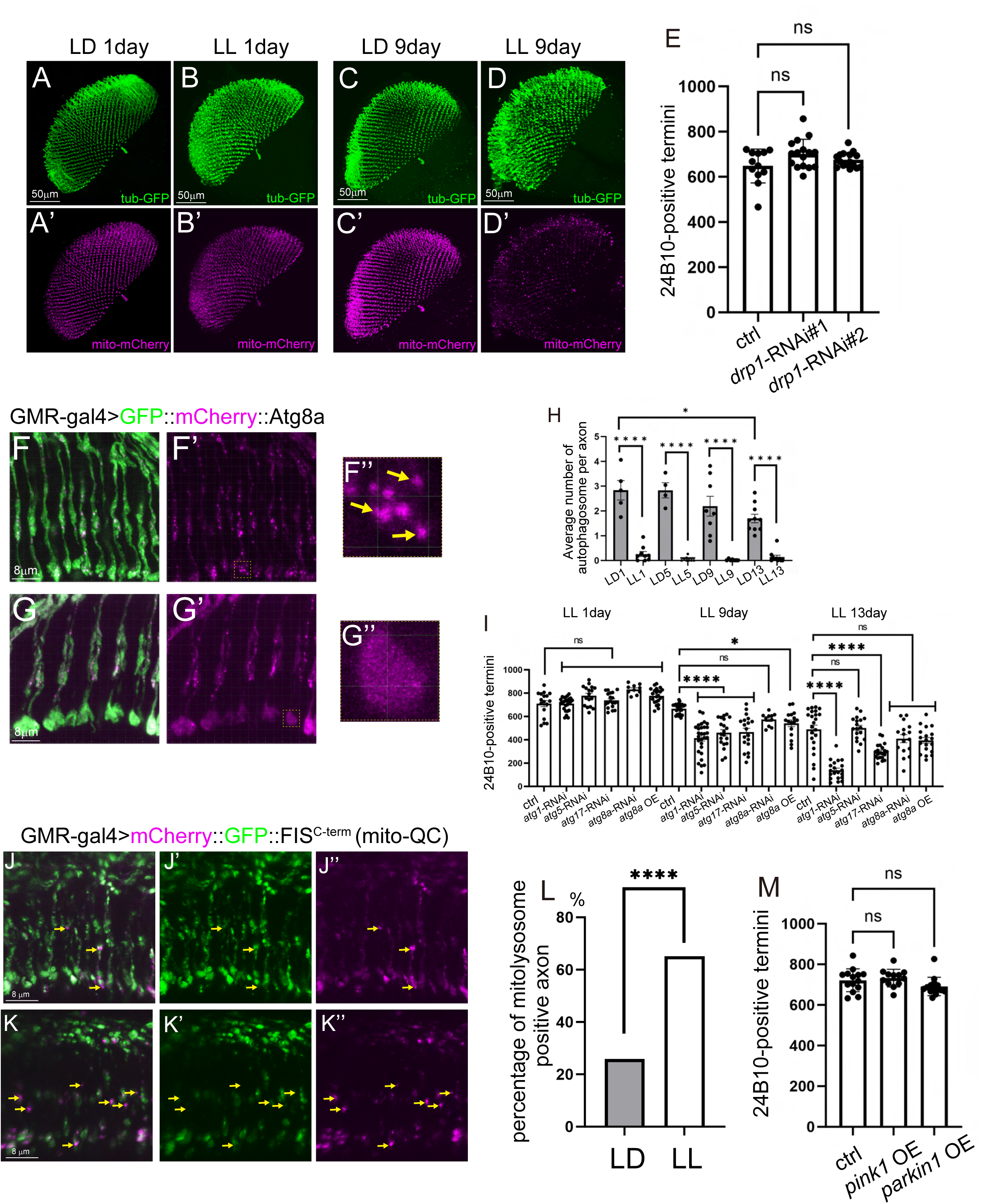
Mitochondrial fragmentation and changes in autophagy and mitophagy under light stress are likely secondary effects. (A–D’) Localization of photoreceptor axons and mitochondria under various light conditions. Photoreceptor axons and mitochondria were visualized using GMR-GAL4 to drive UAS-tub-GFP and UAS-mito-mCherry, respectively. LD_1day (A and A’), LL_1day (B and B’), LD_9day (C and C’), LL_9day (D and D’). Scale bar, 50 μm. (E) Quantification of 24B10-positive photoreceptor axons after 13 days of light stress in control flies and flies expressing *drp1*-RNAi (#1: v44156, #2: BDSC 27682). Each point represents one brain; bars indicate mean ± SD. (F–G’’) Localization of autophagy markers under light stress conditions. Autophagosomes in photoreceptor axons were visualized using UAS-GFP::mCherry::Atg8a, a reporter that detects autolysosomes (mCherry⁺-only puncta) via pH-dependent GFP quenching in acidic lysosomes. 1-day LD (F–F’’) and 1-day LL (G–G’’. Scale bar, 8 μm. Arrows indicate autolysosomes. (H) Quantification of autolysosome number under several light conditions. Each point represents one axon; bars indicate mean ± SD. (I) Quantification of 24B10-positive photoreceptor axons under several light conditions in control and genetically manipulated flies targeting autophagy-related genes. Each point represents one brain; bars indicate mean ± SD. (J–K’) Localization of mitolysosomes under 5 days of LL. Mitolysosomes in photoreceptors were visualized using mito-QC, a reporter composed of a mitochondria-targeted GFP-mCherry tandem tag. Due to GFP quenching in the acidic lysosomal environment, mitolysosomes appear as mCherry⁺-only puncta. Scale bar, 8 μm. Arrows indicate mitolysosomes. (L) Quantification of mitolysosomes under 5 days of LD and LL conditions. (M) Quantification of 24B10-positive photoreceptor axons after 13 days of LL in control flies and flies overexpressing (OE) *pink1* or *parkin*. Each point represents one brain; bars indicate mean ± SD. Statistical treatment of the number of 24B10-positive termini in (E, I, and M) and the average number of autophagosome in (H) were analyzed using the Kruskal–Wallis test with Dunn’s multiple comparisons test. The percentage of the mitolysosome positive axon in (L) was statistically analyzed with the Chi-square test.

**Figure S2.**
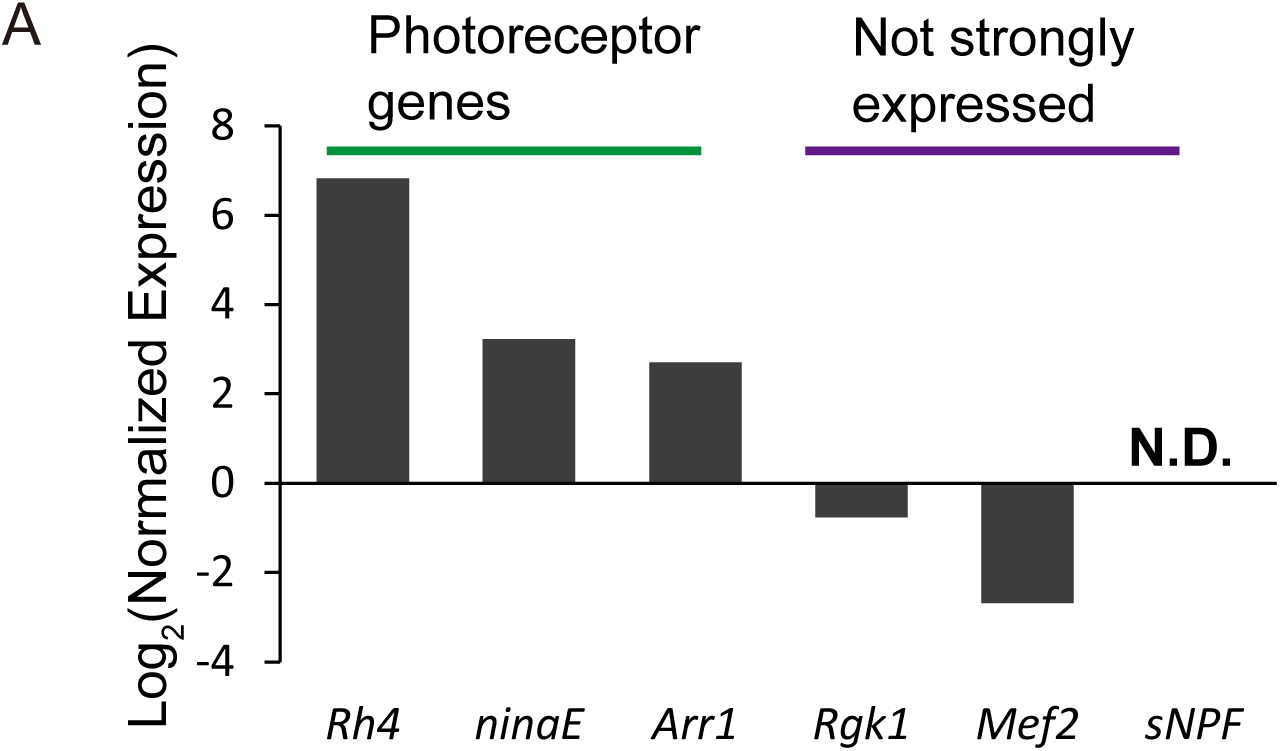
Validation of photoreceptor-specific transcript enrichment by TRAP using RT-qPCR. (A) Quantitative RT-PCR was performed to evaluate the efficiency of TRAP (Translating Ribosome Affinity Purification) in enriching photoreceptor-specific transcripts. Two types of RNA samples were prepared: one before immunoprecipitation (input) and one after ribosome immunoprecipitation (IP-enriched). We analyzed the expression levels of genes known to be highly expressed in photoreceptors (*Rh4*, *ninaE*, *Arr1*), genes with low or absent expression in photoreceptors (*Rgk1*, *Mef2*, *sNPF*), and several housekeeping genes (*ATPsynB*, *αTub84B*, *RP49*, *Ubi-p5E*).^61^ Expression levels were normalized to the housekeeping genes, and enrichment was assessed by comparing transcript abundance before and after TRAP. As expected, photoreceptor-specific genes were highly enriched in the IP samples, while non-photoreceptor genes showed minimal enrichment.

**Figure S3.**
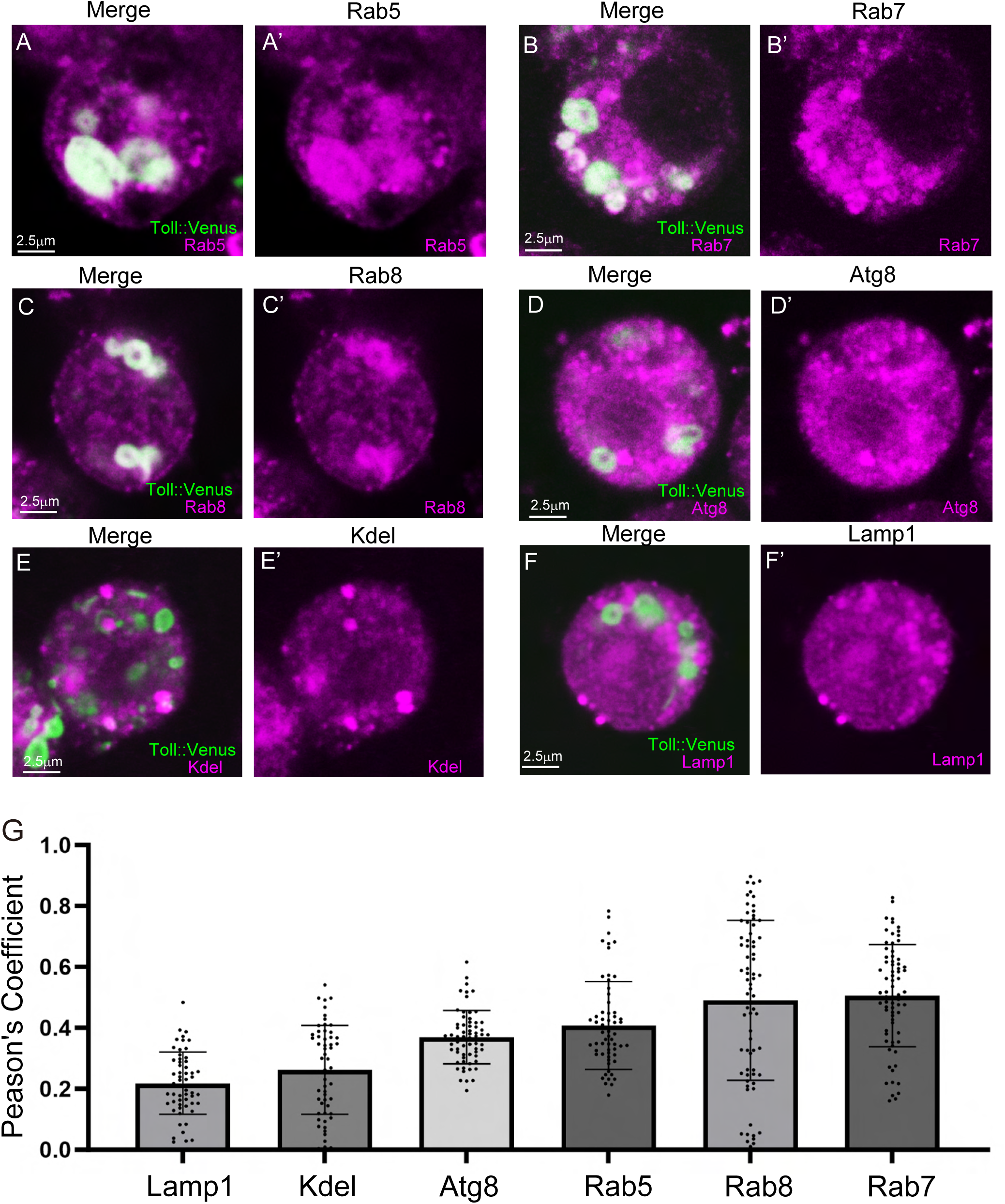
Toll co-localizes with endosomal markers in S2 cells. (A–G’) Co-localization analysis of Toll and subcellular compartment markers in *Drosophila* S2 cells. Toll-1::Venus was expressed by co-transfection of pActin-GAL4 and p20×UAS-Toll-1::Venus plasmids, followed by immunostaining to assess co-localization. (A and A’) Toll-1::Venus anti-Rab5 labeling. (B and B’) Toll-1::Venus anti-Rab7 labeling. (C and C’) Toll-1::Venus anti-Rab8 labeling. (D and D’) Toll-1::Venus anti-Atg8 labeling. (E and E’) Toll-1::Venus anti-Kdel labeling. (F and F’) Toll-1::Venus anti-Lamp1 labeling. Scale bar, 2.5 μm. (G) Quantification of co-localization using Pearson’s correlation coefficients between Toll-1::Venus and various organelle markers: lysosomes (Lamp1), endoplasmic reticulum (Kdel), autophagosomes (Atg8), early endosomes (Rab5), secretory vesicles (Rab8), and late endosomes (Rab7). Each point represents one slice image; bars indicate mean ± SD.

**Figure S4.**
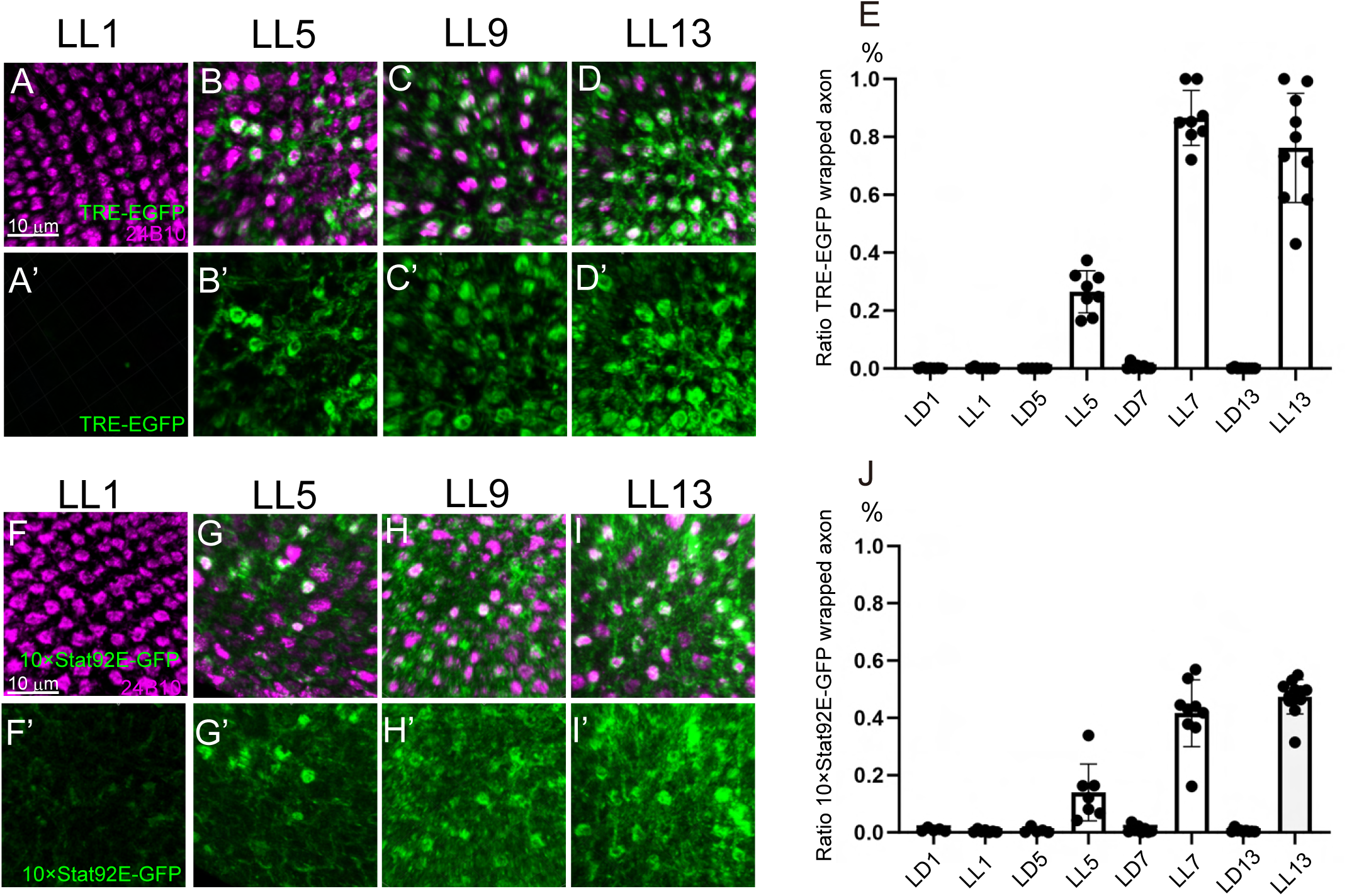
Light stress increases TRE- and Stat92E-dependent transcriptional activity. (A–D’) Time-course of TRE-EGFP reporter signal under chronic light stress. The artificial tetradecanoylphorbol acetate response element (TRE), which contains optimized AP-1 (Jun/Fos) binding sites, is placed upstream of a basal promoter driving EGFP expression. Photoreceptor axons are visualized by 24B10 staining. Scale bar, 2.5 μm (A and A’) Flies after 1 day of LL. Merge (A), TRE-EGFP signal (A’). (B and B’) After 5 days of LL. Merge (B), TRE-EGFP (B’). (C and C’) After 9 days of LL. Merge (C), TRE-EGFP (C’). (D and D’) After 13 days of LL. Merge (D), TRE-EGFP (D’). (E) Quantification of the proportion of photoreceptor axons with TRE-EGFP signal. Each point represents one brain; bars indicate mean ± SD. (F–I’) Stat92E reporter signal using the 10×Stat92E-GFP transgene. This reporter contains Stat92E-binding sequences derived from the *Socs36E* locus and drives EGFP expression under a minimal promoter. Scale bar, 2.5 μm (F and F’) Flies after 1 day of LL. Merge (F), 10×Stat92E-GFP (F’). (G and G’) After 5 days of LL. Merge (G), 10×Stat92E-GFP (G’). (H and H’) After 9 days of LL. Merge (H), 10×Stat92E-GFP (H’). (I and I’) After 13 days of LL. Merge (I), 10×Stat92E-GFP (I’). (J) Quantification of the proportion of photoreceptor axons with 10×Stat92E-GFP signal. Each point represents one brain; bars indicate mean ± SD.

**Figure S5.**
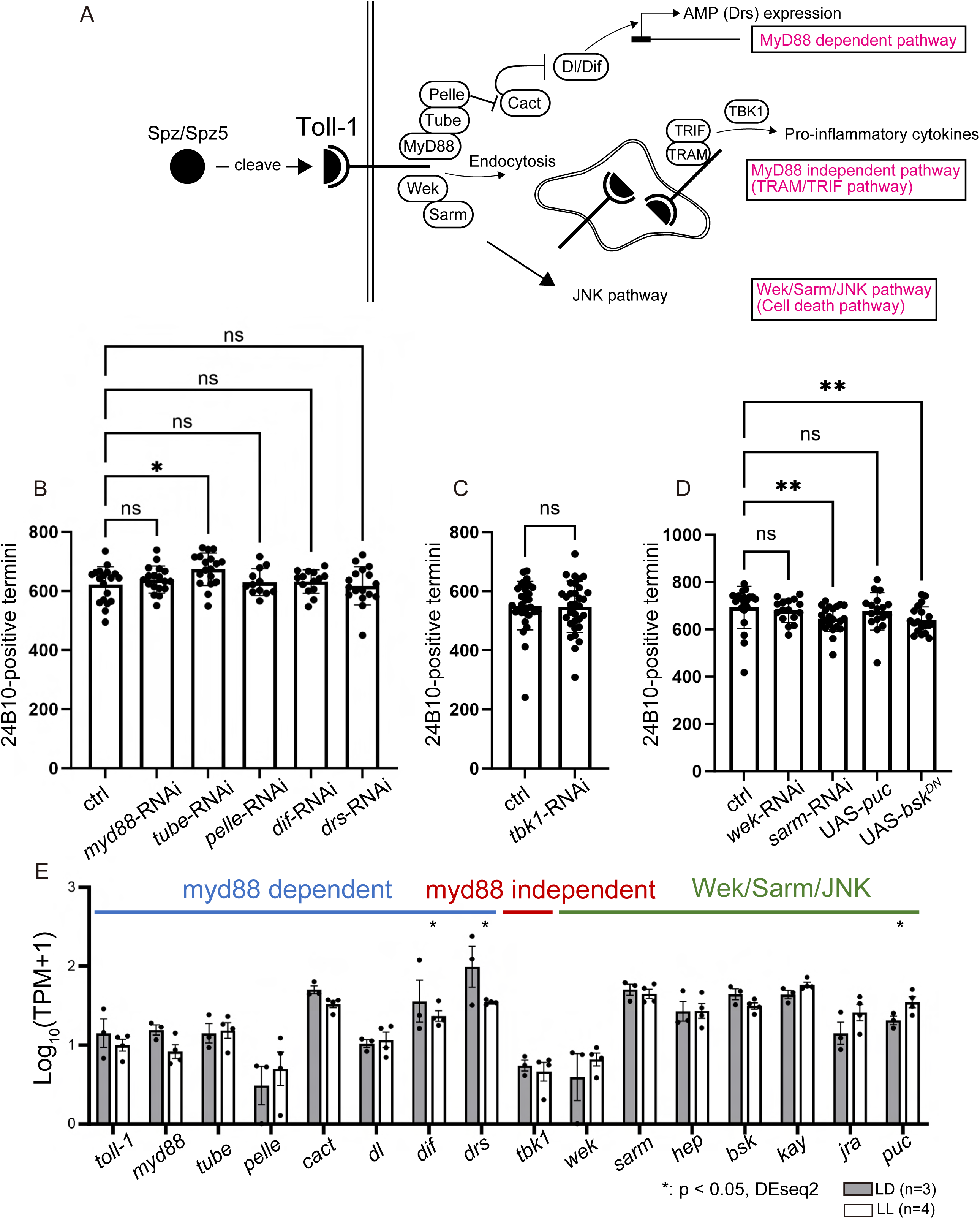
Inhibition of canonical Toll downstream pathways fails to suppress light-induced neurodegeneration. (A) Schematic overview of the Toll-1 signaling pathway. In *Drosophila*, Toll-1 signaling can be divided into three major downstream branches: the MyD88-dependent pathway, which is primarily involved in immunity and embryonic development; the TRAM/TRIF pathway, which is MyD88-independent; and the Wek/Sarm/JNK pathway, which is associated with cell death. (B) Quantification of 24B10-positive photoreceptor axons in flies exposed to 13 days of LL. Comparisons were made between control flies (40D-UAS) and those expressing RNAi constructs targeting MyD88 pathway components specifically in photoreceptors. Each point represents one brain; bars indicate mean ± SD. (C) Quantification of 24B10-positive photoreceptor axons following 13 days of LL flies with RNAi-mediated knockdown of TRAM/TRIF pathway genes in photoreceptors. Among TRAM, TRIF, and TBK1, only TBK1 is conserved in *Drosophila* as the homolog IKKε. Each point represents one brain; bars indicate mean ± SD. (D) Quantification of 24B10-positive photoreceptor axons following 13 days of LL flies with RNAi knockdown of Wek/Sarm/JNK pathway genes in photoreceptors. JNK pathway was suppressed by expressing UAS-*puc* or UAS-*bsk^DN^*, known inhibitors of the pathway. Each point represents one brain; bars indicate mean ± SD. (E) Comparison of expression levels under LD and LL conditions for genes involved in canonical downstream components of the Toll-1 pathway, including the Myd88-dependent pathway, the TRIF/TRAM (Myd88-independent) pathway, and the Wek/Sarm/JNK pathway. Statistical treatment of the number of 24B10-positive termini in (B and D) were analyzed using the Kruskal–Wallis test with Dunn’s multiple comparisons test. The number of 24B10-positive termini in (C) was statistically analyzed with the Mann-Whitney test. Statistical analysis of the expression data in (E) was performed using DESeq.

## SUPPLEMENTALY TABLE

**Table S1, The list of oligonucleotides used in this study.**

**Table S2, The list of genotypes of flies used in this study.**

## Key Resource Table

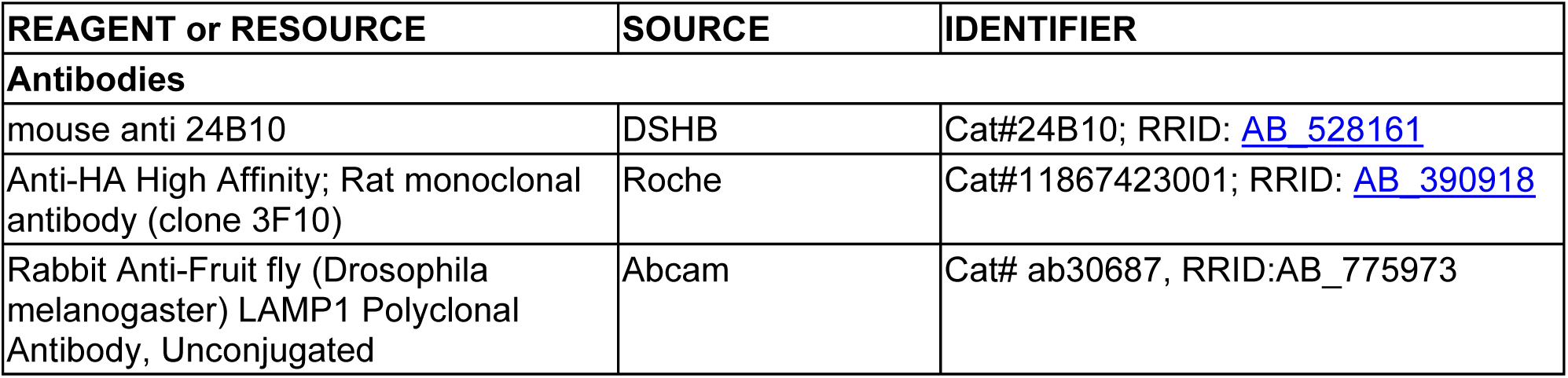
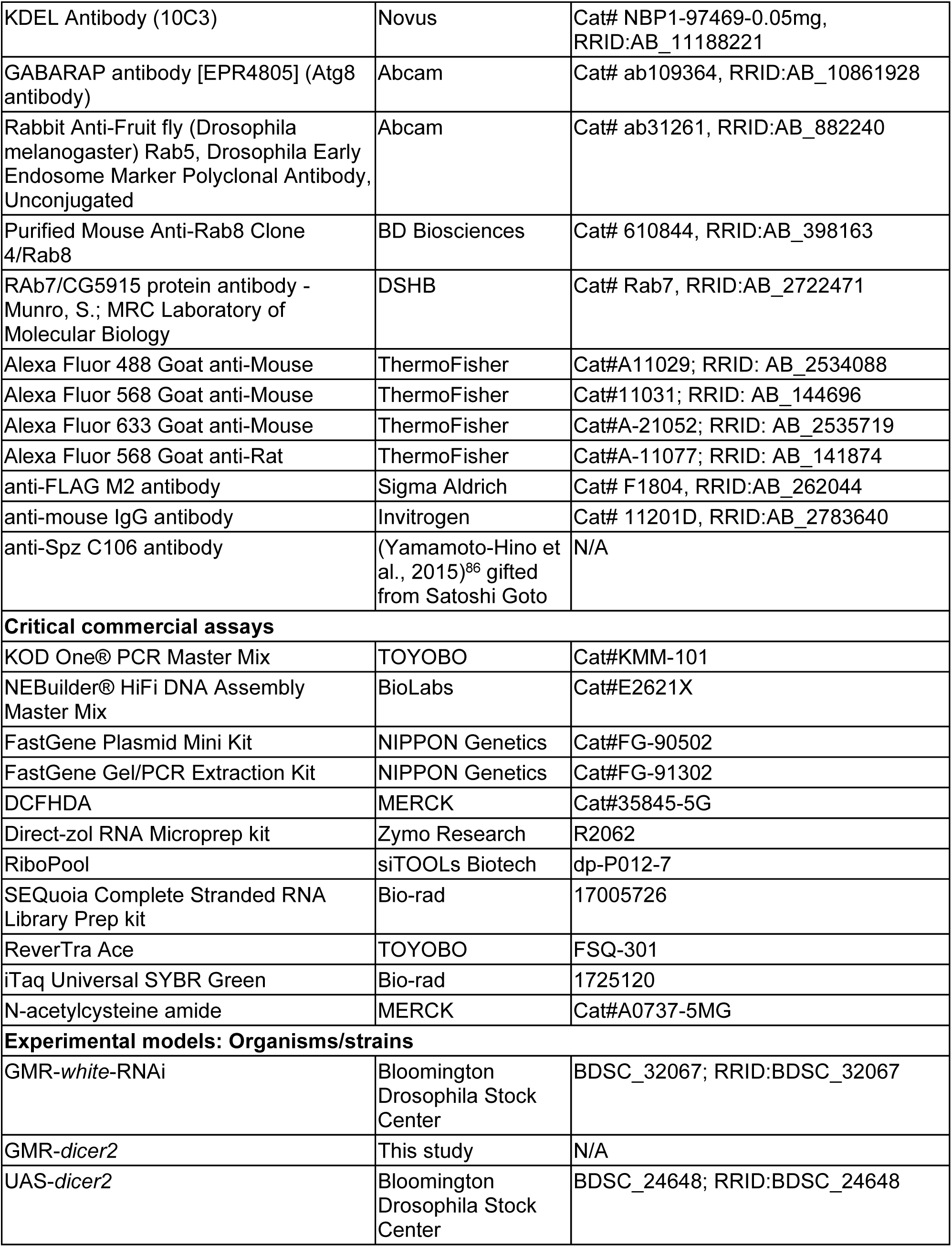
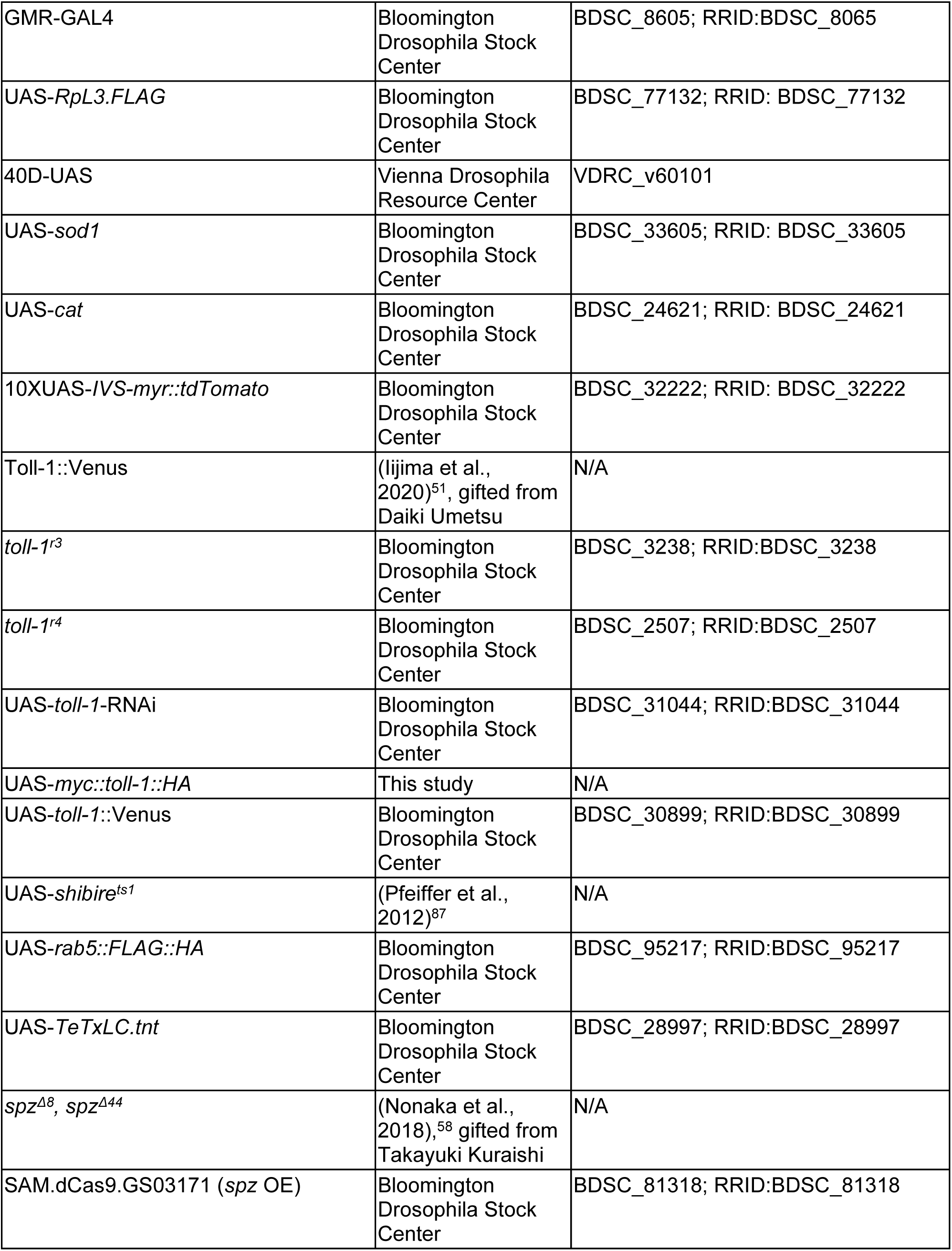
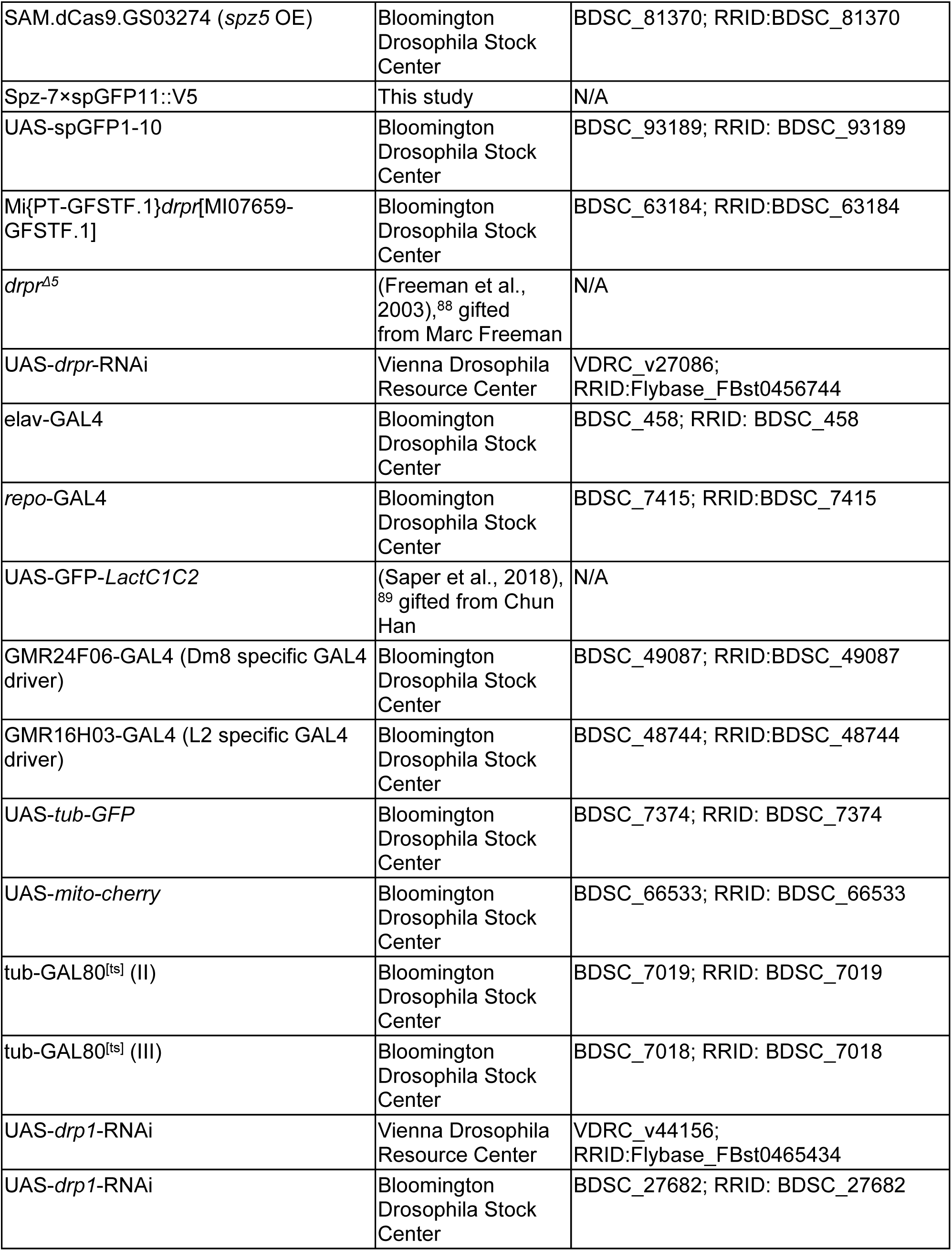
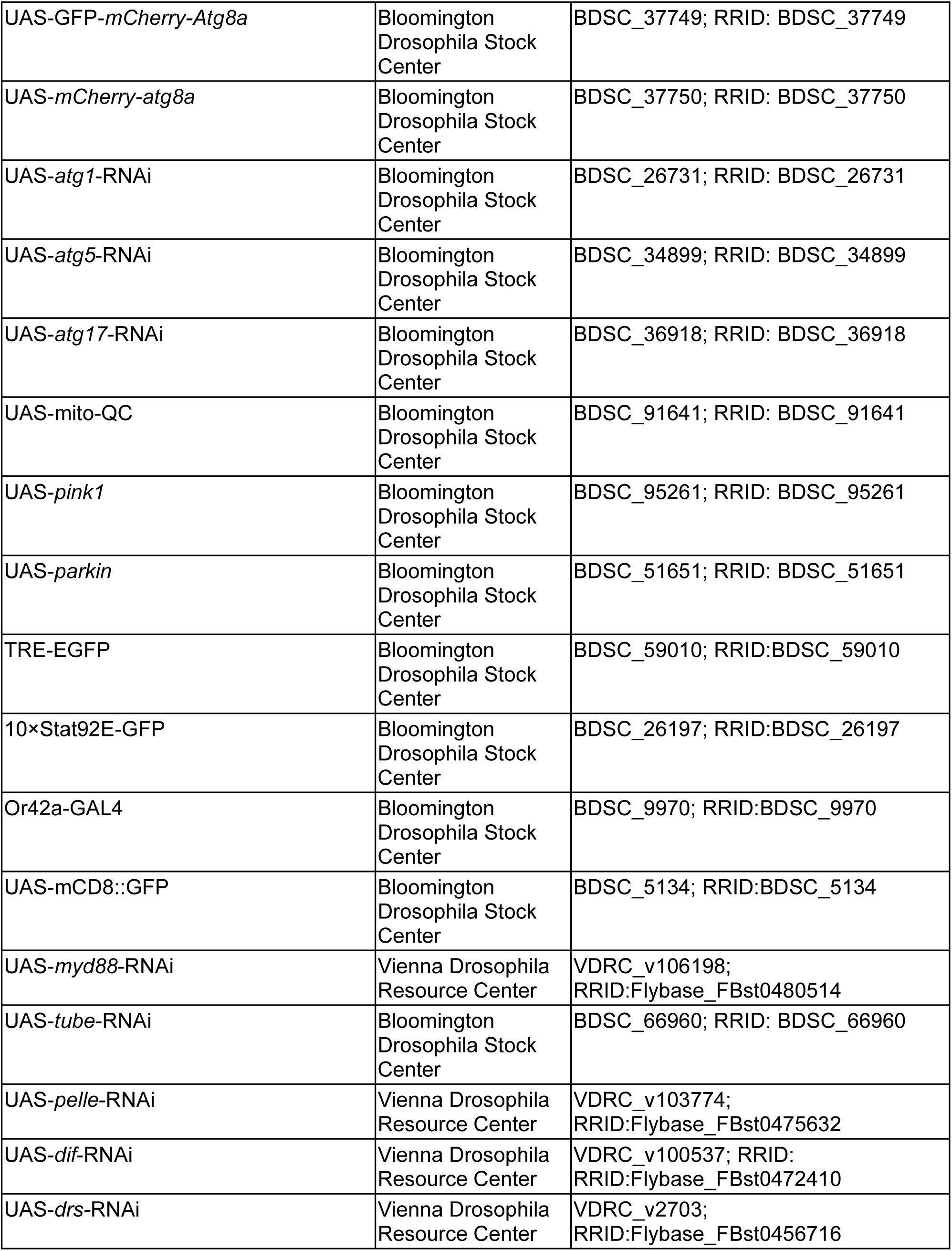
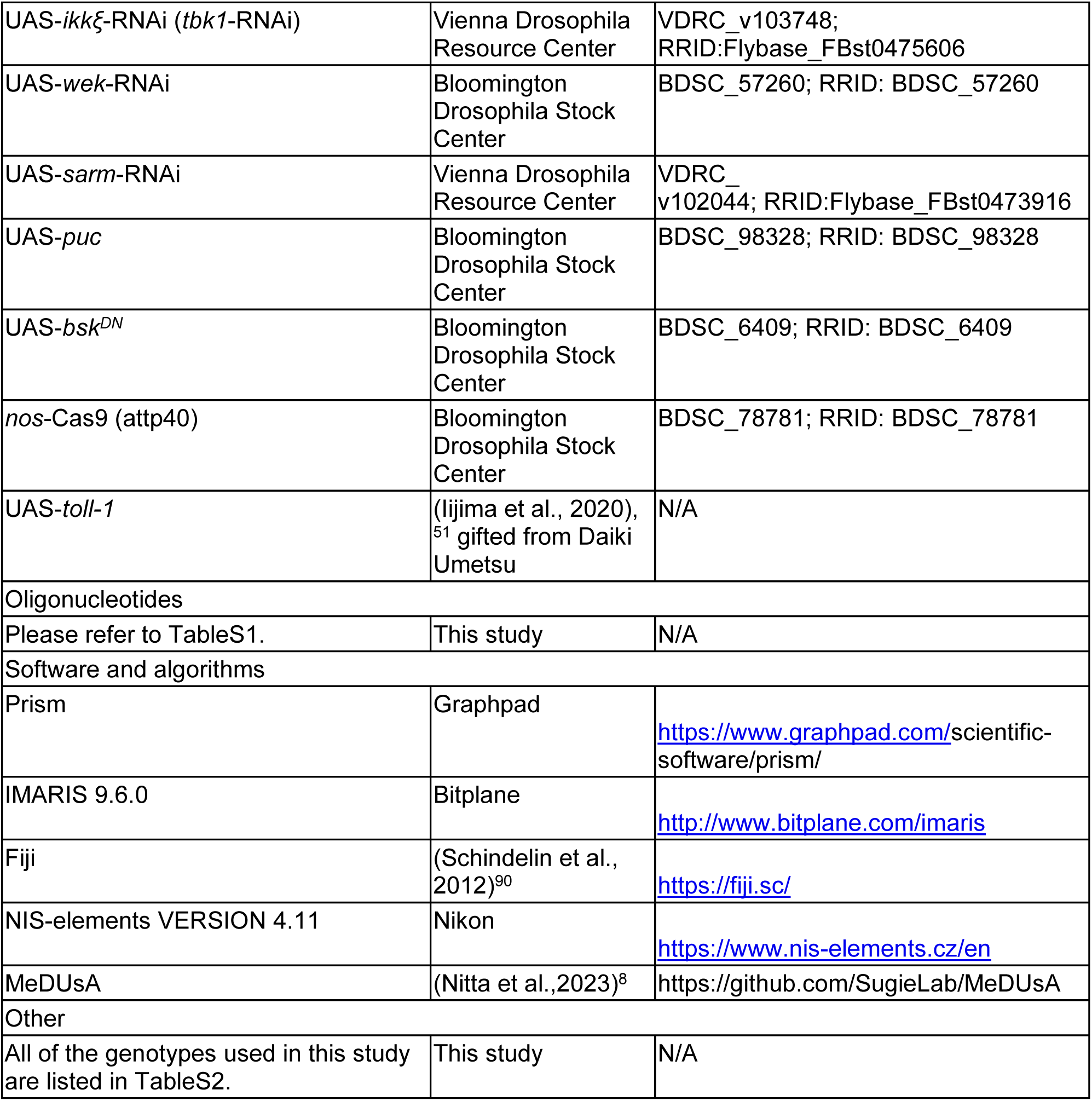

## RESOURCE AVAILABILITY

### Lead Contact

Further information and requests for resources and reagents should be directed to and will be fulfilled by the Lead Contact, Atsushi Sugie.

### Material Availability

All unique reagents generated in this study are available from the Lead Contact without restriction.

### Data and code availability

- All data reported in this paper are available from the Lead Contact upon request.
- This paper does not report original code.
- Any additional information required to reanalyze the data reported in this paper is available from the lead contact upon request.

## EXPERIMENTAL MODEL AND SUBJECT PARTICIPANT DETAILS

### Fly husbandry

Flies were raised in vials with a 2 cm layer of cornmeal and yeast-based food. The ingredients for the fly food and their quantities were as follows: The ingredients for the fly food and their quantities were as follows: water (2 L), agar (23 g), granulated sugar (94 g), glucose (190 g), yeast (101 g, dissolved in 600 mL of water), cornmeal (230 g, dissolved in 700 mL of water), calcium chloride (2.25 g), and potassium bitartrate (27.15 g). First, agar was added to water and heated until boiling. The heat was then reduced, and once the temperature reached 90°C, granulated sugar, glucose, dissolved yeast, dissolved cornmeal, calcium chloride, and potassium bitartrate were added. The mixture was stirred continuously for 15 minutes. After turning off the heat, 6.015 g of Nipagin dissolved in 30 mL of ethanol was added and mixed thoroughly. Finally, when the temperature cooled to 60°C, the prepared medium was dispensed into vials and bottles. For antioxidant treatment, N-acetylcysteine amide (NACA; MERCK, Cat#A0737-5MG) was dissolved in standard fly food at a final concentration of 160 µg/mL. Control flies were maintained on the same food batch without drug supplementation. Flies were maintained at 25 °C under a 12-hour light / 12-hour dark cycle unless otherwise indicated. For experiments under constant light (LL) conditions, an LED lamp (6 W, warm white; Heitronic) was used to illuminate the flies continuously at an intensity of 10,000 lux. The illumination level was measured at the surface of the food medium inside the vial, aligned with the direction of the light source, using a Volt Craft MS-1300 photometer (Conrad Electronic). For EB treatment, black pupae were placed in an approximately 3,860 mL glass chamber containing a 1.5 mL tube filled with 1 mL of 15% EB in mineral oil or 1 mL of mineral oil alone as a control, with the tube cap left open to allow volatilization. Flies were housed in vials covered with a mesh material so that the odor could diffuse into the vials. Newly eclosed adult flies were collected the next day and subsequently maintained in the same chamber under EB or mineral oil exposure for an additional 2 days.

GMR-*white*-RNAi (BDSC 32067), UAS-*dicer2* (BDSC 24648), GMR-GAL4 (BDSC 8605), UAS-*RpL3.FLAG* (BDSC 77132), UAS-*sod1* (BDSC 33605), UAS-*cat* (BDSC 24621), 10XUAS-*IVS-myr::tdTomato* (BDSC 32222), GMR-GAL4 (BDSC 8605), *toll^r3^* (BDSC 3238), *toll^r4^*(BDSC 2507), UAS-*toll-1*-RNAi (BDSC 31044), UAS-*toll-1::Venus* (BDSC 30899), UAS-*rab5::FLAG::HA* (BDSC 95217), UAS-*TeTxLC.tnt* (BDSC 28997), SAM.dCas9.GS03171 (BDSC 81318), SAM.dCas9.GS03274 (BDSC 81370), UAS-spGFP1-10 (BDSC 93189), Mi{PT-GFSTF.1*}drpr*[MI07659-GFSTF.1] (BDSC 63184), elav-GAL4 (BDSC 458), repo-GAL4 (BDSC 7415), GMR24F06-GAL4 (BDSC 49087), GMR16H03-GAL4 (BDSC 48744), UAS-tub-GFP (BDSC 7374), UAS-mito-cherry (BDSC 66533), tub-GAL80ts (II) (BDSC 7019), tub-GAL80ts (III) (BDSC 7018), UAS-*drp1*-RNAi (BSDC 27682), UAS-*GFP-mCherry-Atg8a* (BDSC 37749), UAS*-mCherry-Atg8a* (BDSC 37750), UAS-*atg1*-RNAi (BDSC 26731), UAS-*atg5*-RNAi (BDSC 34899), UAS-*atg17*-RNAi (BSDC 36918), UAS-mito-QC (BSDC 91641), UAS-*pink1* (BDSC 95261), UAS-*parkin* (BDSC 51651), TRE-EGFP (BDSC 59010), 10×Stat92E-GFP (BDSC 26197), Or42a-GAL4 (BSDC 9970), UAS-mCD8::GFP (BSDC 5134), UAS-*tube*-RNAi (BDSC 6690), UAS-*wek*-RNAi (BDSC 57260), UAS-*puc* (BSDC 98328), UAS-*bsk^DN^* (BDSC 6409), and *nos*-Cas9 (attp40) (BDSC 78781), were obtained from Bloomington *Drosophila* Stock Center. 40D-UAS (VDRC 60101), UAS-*drpr*-RNAi (VDRC 27086), UAS-*drp1*-RNAi (VDRC 44156), UAS-*myd88*-RNAi (VDRC 106198), UAS-*pelle*-RNAi (VDRC 103774), UAS-*dif*-RNAi (VDRC 100537), UAS-*ikkξ*-RNAi (VDRC 103748), and UAS-*sarm*-RNAi (VDRC 102044) were obtained from Vienna *Drosophila* Resource Center. Spz-7×spGFP11::V5, 20×UAS-*myc::toll-1::HA* and GMR-dicer2 were newly generated in this study (as described below).

Experiments were conducted at 25 °C unless otherwise noted. For overexpression experiments (Figures 1H, 2F, 3H, 4E–I, and S1M), RNAi-mediated knockdown (Figures 2E, 5P, S1E, and S1I), Shibire^ts1^ inactivation assays (Figure 3D and 3F), TNT experimtnts (Figure 3G), and Toll-1::Venus expressed experiment (Figure 3H), flies were reared at 20 °C until eclosion and then shifted to 29 °C under LL until the day of dissection. For EB exposure experiments (Figure 7B–H), flies were reared at 25 °C and exposed to EB at 23 °C, following experimental conditions described in a previous study.^66^

## METHOD DETAILS

### Generation of transgenic flies

Spz-7×spGFP11::V5 knock-in allele were generated by CRISPR/Cas9 technology. A knock-in vector containing the homology arms, and the cassette (7×spGFP11::V5), was generated as described below. A synthetic DNA construct was designed by fusing a V5 tag to the C-terminus of a sequence containing seven tandem repeats of spGFP11, each separated by a GS linker. Subsequently, the 7×spGFP11::V5 fragment was cloned into a plasmid previously used in earlier studies, which contained an FRT-stop-FRT-GFP cassette, using this plasmid as the template backbone^91^. The accuracy of the inserted sequence was confirmed by DNA sequencing. Approximately 1000 bp fragments flanking the stop codon of Spz were PCR-amplified from the *Drosophila* genome. To avoid cleavage of the knock-in vectors by the two gDNAs, non-synonymous mutations were introduced into the gDNA target sites within these fragments, which were then cloned together with a 7×spGFP11::V5 cassette. The oligo DNAs used for creating the 7×spGFP11::V5 vector, amplification of fragments, and creating gDNAs are listed in **Table S1**. Two gRNA vectors were created and cloned into pBFv-U6.2 vector and were injected into eggs of nos-Cas9 flies at WellGenetics together with the knock-in vector. The precise integration of the knock-in vector was verified by PCR and sequencing.

GMR-*dicer2* and 20×UAS-*myc::toll-1::HA* was generated φC31 integrase-mediated recombination.^92^ Plasmid for generating GMR-*dicer2* was made by Vector Builder by fusing the promoter region of long GMR and dicer2 and injected into eggs of y[1]w[*] P{y[+t7.7]=CaryIP}su(Hw)attP8 at WellGenetics. The cDNA of *toll-1* was amplified from UAS-Toll^FL^-1 flies^51^ by genome extractions. The Toll cDNA was amplified by PCR with a myc tag added to its N-terminus and an HA tag to its C-terminus. The resulting fragment was then cloned into the pGP-20×UAS-IVS-Syn21-jGCaMP8s-p10 vector (Addgene #162386) after digestion with XhoI and XbaI. The primers used to generate the flies are shown in **Table S1**.

### Immunohistochemistry

Brain dissection, fixation, and immunostaining were performed as outlined in the following sentences. Fly brains were dissected in 0.3% PBT (PBS containing Triton X-100) and then fixed with 4% paraformaldehyde at room temperature for 60 min. The samples were washed with 0.3 % PBT 3 times and then a primary antibody was added, followed by incubation overnight at 4 degrees. The next day, the samples were washed with 0.3 % PBT 3 times and then a primary antibody was added, followed by incubation overnight at 4 degrees. Next day, the samples were washed once with 0.3% PBT 3 times. Finally, samples were washed with PBS once and replaced by VECTASHIELD®. For S2 cells, 1×10^5^ of cells were allowed to attach to the glass slide at room temperature for 60 min before the fixation. The following antibodies were used for immunohistochemistry: mAb24B10 (1:25, DSHB), rat antibody to HA (3F10, 1:50, Roche), rabbit antibody to Lamp1 (1:200, Abcam), mouse antibody to KDEL (10C3, 1:50, Novus), rabbit antibody to Atg8 (1:200, Abcam), rabbit antibody to Rab5 (1:50, Abcam), mouse antibody to Rab8 (1:400, BD Biosciences), mouse antibody to Rab7 (1:100, DSHB), and rat antibody to Spz (1:50, C106). The secondary antibodies were Alexa488-, Alexa568- or Alexa633-conjugated (1:400, Life Technologies).

### Image acquisition

All confocal images were taken by Nikon C2+, A1 or Olympus FV3000 confocal microscope. Each brain image was acquired with a Z step size of 0.5 or 1μm. Images were processed with Imaris, Adobe Photoshop, and Adobe Illustrator.

### Automated quantification of photoreceptor axon terminals using MeDUsA

Confocal z-stacks of whole-mount adult fly brains stained with 24B10 were analyzed using MeDUsA (Method for the Quantification of Degeneration Using Fly Axons) to quantify R7 axon terminals in the medulla.^8^ MeDUsA is a Python-based tool that combines a pre-trained deep learning segmentation model with an axon terminal counting algorithm. The software takes the 24B10-labeled confocal stack as input, automatically segments the medulla region and identifies individual R7 photoreceptor axon terminals, then outputs a count of the 24B10-positive R7 axon terminals per brain.

### S2 cell transfection

We used a mixture of culture medium; 500mL of Schneider’s Drosophila Medium (×1, Gibco), 5mL of Penicillin-Streptomycin Solution (×100, Wako), and 50mL of fetal bovine serum (Sigma). S2 cells were grown in 24 well plates (TrueLine) with 500μL of the culture medium. For transfection, 1.5μg of UAS vectors and pActin5C-gal4 were co-transfected with 10μL of HilyMax (DOJINDO) and 30μL of Schneider’s Drosophila Medium and incubated.

The 20×UAS-*toll-1::Venus* plasmids used in S2 cells were generated by cloning Toll cDNA, which was amplified from the 20×UAS-*myc::toll-1::HA* plasmid described in the *Generation of transgenic flies* section. The Toll cDNA was then further amplified by PCR, with the Venus coding sequence added to its C-terminus. All primers used for generating the UAS plasmids are listed in **Table S1**.

### ROS detection and quantification

ROS levels were measured using 10 μM DCFH-DA (Merck), following a procedure adapted from our previous study using MitoSOX-based ROS detection.^93^ Freshly dissected adult fly brains were prepared in PBS and incubated in 10 μM DCFH-DA solution for 15 minutes in the dark at room temperature. After incubation, the brains were sequentially washed with PBS containing 20%, 40%, and 60% Vectashield (Vector Laboratories), then mounted in 100% Vectashield. To quantify ROS levels in photoreceptor R7/8 axons, imaging data were analyzed using IMARIS software (Bitplane). Axon surfaces were automatically generated based on mtdTomato signals driven by GMR-GAL4. The DCF channel was then masked using the mtdTomato-defined surfaces. ROS levels were calculated as the ratio of DCF-positive voxel volume to the total mtdTomato surface voxel volume. All quantifications were performed in a blinded manner with respect to genotype.

### Characterization of Toll-1::Venus puncta in axons

Analysis of Toll-1::Venus puncta was performed manually using the Spot function in Imaris software (Bitplane). The region visualized by the anti-24B10 antibody was defined as the R7/8 axonal area. Within this region, punctate Toll-1::Venus signals were manually marked using the Spot function. The number of Toll-1::Venus puncta per axon was calculated. All quantifications were performed in a genotype-blinded manner.

### Quantification of Spz signal

Reconstituted GFP signal from endogenous Spz was quantified relative to 24B10-labeled photoreceptor axons using IMARIS software (Bitplane). A surface corresponding to the 24B10-positive photoreceptor axon region was first generated using the Surface function in Imaris. Within this 24B10-defined region, reconstructed Spz signal was segmented again using the Surface function, and its voxel volume was measured. The ratio of Spz voxel volume to 24B10 voxel volume was then calculated for each sample. All quantifications were performed in a genotype-blinded manner.

### Analysis of axon terminal wrapping by Drpr::GFP or GFP-Lact^C1C2^

Quantification was performed using Imaris software (Bitplane). The region labeled with the anti-24B10 antibody was defined as the R7/8 axonal area. Within this region, R7 axon terminals in the M4–M6 layers were identified based on the 24B10 signal. Axons showing clear accumulation of Drpr::GFP or GFP-LactC1C2 at their terminals were defined as “wrapped axons.” The percentage of wrapped axons was calculated by dividing the number of wrapped axons by the total number of axons. All quantifications were conducted in a genotype-blinded manner.

### Co-localization analysis Toll-1::Venus and antibodies

Co-localization analysis was performed using the JACoP plugin in Fiji (ImageJ).^90,94^ Single optical slices capturing the central region of cultured cells expressing Toll-1::Venus were acquired with a confocal microscope. Pearson’s correlation coefficient was calculated to assess the degree of co-localization between Toll-1::Venus and antibody signals.

### Analysis of autophagosome in photoreceptor axons

Autophagosome formation in the R7/8 photoreceptor axons of the optic lobe was analyzed using the tandem fluorescent-tagged autophagy reporter UAS-GFP-mCherry-Atg8a, expressed under the control of GMR-GAL4. This reporter visualizes autolysosomes (mCherry⁺-only puncta) based on the pH-sensitive quenching of GFP in acidic lysosomal compartments. Manual quantification was performed using the Spot function in Imaris software (Bitplane). Puncta were identified based on the fluorescent signals of GFP-mCherry-Atg8a in R7/8 axons, and only mCherry⁺-only puncta were counted as autolysosomes. The average number of mCherry⁺-only puncta per axon within each optic lobe was quantified. All quantifications were carried out in a genotype-blinded manner.

### Mitolysosome analysis in photoreceptor axons

Mitophagy in the R7/8 photoreceptor axons of the medulla was assessed using the mito-QC reporter, which consists of a mitochondrial-targeted tandem GFP-mCherry tag. Due to the quenching of GFP in the acidic environment of lysosomes, mitolysosomes can be identified as mCherry⁺-only puncta. These red-only puncta were manually annotated using the Spot function in Imaris software (Bitplane). The average number of mitolysosomes per axon was calculated for each medulla by dividing the total number of mCherry⁺-only puncta by the estimated number of axons. All quantifications were performed in a genotype-blinded manner.

### Analysis of axon terminal wrapping by TRE-EGFP or 10×Stat92E-GFP

To analyze axon terminal wrapping by TRE-EGFP or 10×Stat92E-GFP, axon terminal regions in the optic lobe were first defined using the 24B10 signal. Axon terminal regions in the optic lobe were delineated from the 24B10 signal using MeDUsA’s automated image analysis pipeline. In this process, MeDUsA generates RGB masks by integrating three z-sections into a color composite, which facilitates identification of axon terminal regions via deep-learning segmentation.^8^ The extracted RGB mask was then integrated into the original Imaris files as a new channel. The surface function of Imaris (Bitplane) was applied to the RGB mask channel to generate the axon terminal surface region. Within this defined surface, objects positive for 24B10 and TRE-EGFP or 10×Stat92E-GFP signals were segmented, and the number of objects in each channel was quantified. The percentage of wrapped axons was calculated by dividing the number of TRE-EGFP or 10×Stat92E-GFP-positive objects by the number of 24B10-positive objects. All quantifications were performed in a blinded manner with respect to genotype.

### Analysis of Or42a ORN volume

Or42a ORN volume was quantified using Imaris software (Bitplane). Or42a ORNs labeled by Or42a-GAL4-driven UAS-mCD8::GFP were reconstructed in each antennal lobe using the Surface function, and the enclosed volume was measured. All quantifications were performed in a blinded manner with respect to genotype.

### Library preparation for TRAP-seq

5- to 12-day-old female flies were flash-frozen with liquid nitrogen, thoroughly vortexed, and the heads were isolated from the bodies with metal mesh in a similar manner reported previously.^46,95^ Approximately 200 frozen heads were mixed with 250 µl of frozen droplets of lysis buffer [20 mM Tris-HCl pH7.5, 150mM NaCl, 5 mM MgCl_2_, 1 mM dithiothreitol, 1 % Triton X-100, 100 µg/ml chloramphenicol, and 100 µg/ml cycloheximide] and two tungsten bullets (5.0 mm diameter, TC50-0020, Bio Medical Science) in a pre-chilled container, then pulverized with grinding at 30 Hz for 30 seconds using TissueLyser II (QIAGEN). Supernatant was recovered after centrifugation at 20,000 g for 10 minutes. RNA concentration was measured with Qubit RNA High-sensitivity kit (Q32852, Thermo Fisher Scientific) and set to be 50 ng/µl with dilution with the lysis buffer.

RNA was purified using TRIzol-LS (10296010, ThermoFisher Scientific) and Direct-zol RNA Microprep kit (R2062, Zymo Research). Ribosomal RNA was depleted using RiboPool (dp-P012-7, siTOOLs Biotech). The libraries were constructed using SEQuoia Complete Stranded RNA Library Prep kit (17005726, Bio-rad) according to the manufacturer’s instruction. The libraries were sequenced with NovaSeq X Plus (150 base pair, paired end, 5 Gb/sample) in NIPPON GENE Co.

### Data analysis

Data were analyzed using the NIG supercomputer at ROIS National Institute of Genetics. Adaptor sequences were removed using Fastp,^96^ and the reads that matched to the non-coding RNA were discarded. The remaining reads were mapped onto the *Drosophila melanogaster* release 6 genome. Mapping was performed using STAR.^97^ The number of mapped reads is as follows:

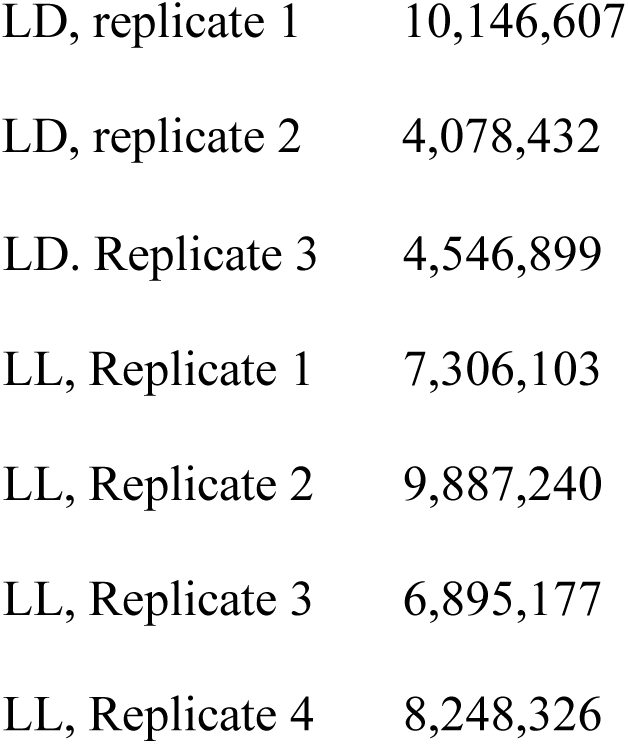

For genes with alternatively spliced transcripts, the isoform with the highest TPM in the LD condition was selected as the ‘representative’ isoform and were considered. P-values were calculated with an *R* package, DEseq2. GO-term enrichment analysis was performed using the Database for Annotation, Visualization, and Integrated Discovery (DAVID).^98,99^

### qRT-PCR

RNA from immunoprecipitated or the whole head samples were purified using TRIzol-LS and Direct-zol RNA Microprep kit as described above. After measuring the RNA concentration, the concentration was made consistent among samples (3.15 ng/ul, 6 ul) and the samples were reverse-transcribed with ReverTra Ace (FSQ-301, TOYOBO). Real-time PCR was performed on cDNA using iTaq Universal SYBR Green (1725120, Bio-rad). Used primer sequences are listed in **Table S1.**

Relative mRNA level of the photoreceptor neuron or not strongly expressed genes were quantified by normalizing by the house-keeping genes (*αtub84B, Rp49, ATPsynb, Ubi-p5E*). Enrichment of these mRNA was calculated as the relative mRNA level in the immunoprecipitated samples divided by the whole head samples (Figure S2).

### Immunoprecipitation

Anti-FLAG M2 antibody (F1804, Sigma Aldrich) and Dynabeads M-280 bound to anti-mouse IgG antibody (11201D, Invitrogen) were used for immunoprecipitation. 10 µl of the beads solution, washed twice with the aforementioned lysis buffer, was mixed with 1 µl of the M2 antibody, and incubated at 4 °C for 1 hour with rotation. Beads were incubated with the 200 ul of lysate at 4 °C for 1 hour with rotation and washed for four times with the lysis buffer. The ribosome-bound mRNA was eluted with 50 µl of 100 µg/ml 3× FLAG peptide (GEN-3XFLAG-25, Protein Ark) dissolved in the lysis buffer. Recovery rate of RNA was around 0.7∼0.9% of the input sample.

### Quantification and statistical analysis

All of the statistical analyses are described in the figure legends. Statistical significance. n.s. *P* > 0.05, * *P* ≤ 0.05, ** *P* ≤ 0.01, *** *P* ≤ 0.001, **** *P* ≤ 0.0001.

