## Supplementary material for "An Innate Immune Receptor Toll-1 converts chronic light stress into glial-phagocytosis": Key Resource Table

### Key resources table

| REAGENT or RESOURCE | SOURCE | IDENTIFIER |
| --- | --- | --- |
| Antibodies | | |
| mouse anti 24B10 | DSHB | Cat#24B10; RRID: [AB_528161](http://antibodyregistry.org/AB_528161" \t "_blank) |
| Anti-HA High Affinity; Rat monoclonal antibody (clone 3F10) | Roche | Cat#11867423001; RRID: [AB_390918](http://antibodyregistry.org/AB_390918" \t "_blank) |
| Rabbit Anti-Fruit fly (Drosophila melanogaster) LAMP1 Polyclonal Antibody, Unconjugated | Abcam | Cat# ab30687, RRID:AB_775973 |
| KDEL Antibody (10C3) | Novus | Cat# NBP1-97469-0.05mg, RRID:AB_11188221 |
| GABARAP antibody [EPR4805] (Atg8 antibody) | Abcam | Cat# ab109364, RRID:AB_10861928 |
| Rabbit Anti-Fruit fly (Drosophila melanogaster) Rab5, Drosophila Early Endosome Marker Polyclonal Antibody, Unconjugated | Abcam | Cat# ab31261, RRID:AB_882240 |
| Purified Mouse Anti-Rab8 Clone 4/Rab8 | BD Biosciences | Cat# 610844, RRID:AB_398163 |
| RAb7/CG5915 protein antibody - Munro, S.; MRC Laboratory of Molecular Biology | DSHB | Cat# Rab7, RRID:AB_2722471 |
| Alexa Fluor 488 Goat anti-Mouse | ThermoFisher | Cat#A11029; RRID: AB_2534088 |
| Alexa Fluor 568 Goat anti-Mouse | ThermoFisher | Cat#11031; RRID: AB_144696 |
| Alexa Fluor 633 Goat anti-Mouse | ThermoFisher | Cat#A-21052; RRID: AB_2535719 |
| Alexa Fluor 568 Goat anti-Rat | ThermoFisher | Cat#A-11077; RRID: AB_141874 |
| anti-FLAG M2 antibody | Sigma Aldrich | Cat# F1804, RRID:AB_262044 |
| anti-mouse IgG antibody | Invitrogen | Cat# 11201D, RRID:AB_2783640 |
| anti-Spz C106 antibody | (Yamamoto-Hino et al., 2015) gifted from Satoshi Goto | N/A |
| Critical commercial assays | | |
| KOD One® PCR Master Mix | TOYOBO | Cat#KMM-101 |
| NEBuilder® HiFi DNA Assembly Master Mix | BioLabs | Cat#E2621X |
| FastGene Plasmid Mini Kit | NIPPON Genetics | Cat#FG-90502 |
| FastGene Gel/PCR Extraction Kit | NIPPON Genetics | Cat#FG-91302 |
| DCFHDA | MERCK | Cat#35845-5G |
| Direct-zol RNA Microprep kit | Zymo Research | R2062 |
| RiboPool | siTOOLs Biotech | dp-P012-7 |
| SEQuoia Complete Stranded RNA Library Prep kit | Bio-rad | 17005726 |
| ReverTra Ace | TOYOBO | FSQ-301 |
| iTaq Universal SYBR Green | Bio-rad | 1725120 |
| N-acetylcysteine amide | MERCK | Cat#A0737-5MG |
| Experimental models: Organisms/strains | | |
| GMR-*white*-RNAi | Bloomington Drosophila Stock Center | BDSC_32067; RRID:BDSC_32067 |
| GMR-*dicer2* | This study | N/A |
| UAS-*dicer2* | Bloomington Drosophila Stock Center | BDSC_24648; RRID:BDSC_24648 |
| GMR-GAL4 | Bloomington Drosophila Stock Center | BDSC_8605; RRID:BDSC_8065 |
| UAS-*RpL3.FLAG* | Bloomington Drosophila Stock Center | BDSC_77132; RRID: BDSC_77132 |
| 40D-UAS | Vienna Drosophila Resource Center | VDRC_v60101 |
| UAS-*sod1* | Bloomington Drosophila Stock Center | BDSC_33605; RRID: BDSC_33605 |
| UAS-*cat* | Bloomington Drosophila Stock Center | BDSC_24621; RRID: BDSC_24621 |
| 10XUAS-*IVS-myr::tdTomato* | Bloomington Drosophila Stock Center | BDSC_32222; RRID: BDSC_32222 |
| Toll-1::Venus | (Iijima et al., 2020), gifted from Daiki Umetsu | N/A |
| *toll-1^r3^* | Bloomington Drosophila Stock Center | BDSC_3238; RRID:BDSC_3238 |
| *toll-1^r4^* | Bloomington Drosophila Stock Center | BDSC_2507; RRID:BDSC_2507 |
| UAS-*toll-1*-RNAi | Bloomington Drosophila Stock Center | BDSC_31044; RRID:BDSC_31044 |
| UAS-*myc::toll-1::HA* | This study | N/A |
| UAS-*toll-1*::Venus | Bloomington Drosophila Stock Center | BDSC_30899; RRID:BDSC_30899 |
| UAS-*shibire^ts1^* | (Pfeiffer et al., 2012) | N/A |
| UAS-*rab5::FLAG::HA* | Bloomington Drosophila Stock Center | BDSC_95217; RRID:BDSC_95217 |
| UAS-*TeTxLC.tnt* | Bloomington Drosophila Stock Center | BDSC_28997; RRID:BDSC_28997 |
| *spz^Δ8^, spz^Δ44^* | (Nonaka et al., 2018), gifted from Takayuki Kuraishi | N/A |
| SAM.dCas9.GS03171 (*spz* OE) | Bloomington Drosophila Stock Center | BDSC_81318; RRID:BDSC_81318 |
| SAM.dCas9.GS03274 (*spz5* OE) | Bloomington Drosophila Stock Center | BDSC_81370; RRID:BDSC_81370 |
| Spz-7×spGFP11::V5 | This study | N/A |
| UAS-spGFP1-10 | Bloomington Drosophila Stock Center | BDSC_93189; RRID: BDSC_93189 |
| Mi{PT-GFSTF.1}*drpr*[MI07659-GFSTF.1] | Bloomington Drosophila Stock Center | BDSC_63184; RRID:BDSC_63184 |
| *drpr^Δ5^* | (Freeman et al., 2003), gifted from Marc Freeman | N/A |
| UAS-*drpr*-RNAi | Vienna Drosophila Resource Center | VDRC_v27086; RRID:Flybase_FBst0456744 |
| elav-GAL4 | Bloomington Drosophila Stock Center | BDSC_458; RRID: BDSC_458 |
| *repo*-GAL4 | Bloomington Drosophila Stock Center | BDSC_7415; RRID:BDSC_7415 |
| UAS-GFP-*LactC1C2* | (Saper et al., 2018), gifted from Chun Han | N/A |
| GMR24F06-GAL4 (Dm8 specific GAL4 driver) | Bloomington Drosophila Stock Center | BDSC_49087; RRID:BDSC_49087 |
| GMR16H03-GAL4 (L2 specific GAL4 driver) | Bloomington Drosophila Stock Center | BDSC_48744; RRID:BDSC_48744 |
| UAS-*tub-GFP* | Bloomington Drosophila Stock Center | BDSC_7374; RRID: BDSC_7374 |
| UAS-*mito-cherry* | Bloomington Drosophila Stock Center | BDSC_66533; RRID: BDSC_66533 |
| tub-GAL80^[ts]^ (II) | Bloomington Drosophila Stock Center | BDSC_7019; RRID: BDSC_7019 |
| tub-GAL80^[ts]^ (III) | Bloomington Drosophila Stock Center | BDSC_7018; RRID: BDSC_7018 |
| UAS-*drp1*-RNAi | Vienna Drosophila Resource Center | VDRC_v44156; RRID:Flybase_FBst0465434 |
| UAS-*drp1*-RNAi | Bloomington Drosophila Stock Center | BDSC_27682; RRID: BDSC_27682 |
| UAS-GFP-*mCherry-Atg8a* | Bloomington Drosophila Stock Center | BDSC_37749; RRID: BDSC_37749 |
| UAS-*mCherry-atg8a* | Bloomington Drosophila Stock Center | BDSC_37750; RRID: BDSC_37750 |
| UAS-*atg1*-RNAi | Bloomington Drosophila Stock Center | BDSC_26731; RRID: BDSC_26731 |
| UAS-*atg5*-RNAi | Bloomington Drosophila Stock Center | BDSC_34899; RRID: BDSC_34899 |
| UAS-*atg17*-RNAi | Bloomington Drosophila Stock Center | BDSC_36918; RRID: BDSC_36918 |
| UAS-mito-QC | Bloomington Drosophila Stock Center | BDSC_91641; RRID: BDSC_91641 |
| UAS-*pink1* | Bloomington Drosophila Stock Center | BDSC_95261; RRID: BDSC_95261 |
| UAS-*parkin* | Bloomington Drosophila Stock Center | BDSC_51651; RRID: BDSC_51651 |
| TRE-EGFP | Bloomington Drosophila Stock Center | BDSC_59010; RRID:BDSC_59010 |
| 10×Stat92E-GFP | Bloomington Drosophila Stock Center | BDSC_26197; RRID:BDSC_26197 |
| Or42a-GAL4 | Bloomington Drosophila Stock Center | BDSC_9970; RRID:BDSC_9970 |
| UAS-mCD8::GFP | Bloomington Drosophila Stock Center | BDSC_5134; RRID:BDSC_5134 |
| UAS-*myd88*-RNAi | Vienna Drosophila Resource Center | VDRC_v106198; RRID:Flybase_FBst0480514 |
| UAS-*tube*-RNAi | Bloomington Drosophila Stock Center | BDSC_66960; RRID: BDSC_66960 |
| UAS-*pelle*-RNAi | Vienna Drosophila Resource Center | VDRC_v103774; RRID:Flybase_FBst0475632 |
| UAS-*dif*-RNAi | Vienna Drosophila Resource Center | VDRC_v100537; RRID: RRID:Flybase_FBst0472410 |
| UAS-*drs-*RNAi | Vienna Drosophila Resource Center | VDRC_v2703; RRID:Flybase_FBst0456716 |
| UAS-*ikkξ*-RNAi (*tbk1*-RNAi) | Vienna Drosophila Resource Center | VDRC_v103748; RRID:Flybase_FBst0475606 |
| UAS-*wek*-RNAi | Bloomington Drosophila Stock Center | BDSC_57260; RRID: BDSC_57260 |
| UAS-*sarm*-RNAi | Vienna Drosophila Resource Center | VDRC_ v102044; RRID:Flybase_FBst0473916 |
| UAS-*puc* | Bloomington Drosophila Stock Center | BDSC_98328; RRID: BDSC_98328 |
| UAS-*bsk^DN^* | Bloomington Drosophila Stock Center | BDSC_6409; RRID: BDSC_6409 |
| *nos*-Cas9 (attp40) | Bloomington Drosophila Stock Center | BDSC_78781; RRID: BDSC_78781 |
| UAS-*toll-1* | (Iijima et al., 2020), gifted from Daiki Umetsu | N/A |
| Oligonucleotides | | |
| Please refer to TableS1. | This study | N/A |
| Software and algorithms | | |
| Prism | Graphpad | [https://www.graphpad.com/](https://www.graphpad.com/" \t "_blank)scientific-software/prism/ |
| IMARIS 9.6.0 | Bitplane | [http://www.bitplane.com/imaris](http://www.bitplane.com/imaris" \t "_blank) |
| Fiji | (Schindelin et al.,) | [https://fiji.sc/](https://fiji.sc/" \t "_blank) |
| NIS-elements VERSION 4.11 | Nikon | [https://www.nis-elements.cz/en](https://www.nis-elements.cz/en" \t "_blank) |
| MeDUsA | (Nitta et al.,) | https://github.com/SugieLab/MeDUsA |
| Other | | |
| All of the genotypes used in this study are listed in TableS2. | This study | N/A |
